# Evaluating institutional open access performance: Methodology, challenges and assessment

**DOI:** 10.1101/2020.03.19.998336

**Authors:** Chun-Kai Huang, Cameron Neylon, Richard Hosking, Lucy Montgomery, Katie Wilson, Alkim Ozaygen, Chloe Brookes-Kenworthy

## Abstract

Open Access to research outputs is becoming rapidly more important to the global research community and society. Changes are driven by funder mandates, institutional policy, grass-roots advocacy and culture change. It has been challenging to provide a robust, transparent and updateable analysis of progress towards open access that can inform these interventions, particularly at the institutional level. Here we propose a minimum reporting standard and present a large-scale analysis of open access progress across 1,207 institutions world-wide that shows substantial progress being made. The analysis detects responses that coincide with policy and funding interventions. Among the striking results are the high performance of Latin American and African universities, particularly for gold open access, whereas overall open access levels in Europe and North America are driven by repository-mediated access. We present a top-100 of global universities with the world’s leading institutions achieving around 80% open access for 2017 publications.

## 1 Introduction

Open access is a policy aspiration for research funders, organisations, and communities globally. While there is substantial disagreement on the best route to achieve open access, the idea that wider availability of research outputs should be a goal is broadly shared. Over the past decade, there has been massive increase in the volume of publications available open access. Piwowar et al. (2018) showed that the global proportion of open access articles was about 45% for those published in 2015, compared to around 5% before 1990. A more recent projection suggests that 44% of all outputs ever published will be freely accessible in 2025 (Piwowar et al., 2019).

This massive increase has been driven in large part by policy initiatives. Medical research funders such as the Wellcome Trust and Medical Research Council in the UK and the National Institutes of Health in the US led a wide range of funder policy interventions. Universities such as Harvard, Liege, Southampton and others developed local polices and infrastructures that became more widely adopted. Plan S, led by a coalition of funders^1^, announced in 2019 has as its goal the complete conversion of scholarly publishing to immediate open access. This is the most ambitious, and therefore the most controversial, policy initiative to date with questions raised about the approach (Rabesandratana, 2019; Haug, 2019; Barbour and Nicholls, 2019), implementation details (McNutt, 2019; Gómez-Fernández,2019; Brainard, 2019; Agustini and Berk, 2019), and unintended side effects for existing programs outside North America and Northwestern Europe (Debat and Babini, 2019; Aguado-López and Becerril-García, 2019).

Despite the scale and success (at least in some areas) of these interventions, there is limited comparative and quantitative research about which policy interventions have been the most successful. In part this is due to a historical lack of high-quality data on open access, the heterogeneous nature of the global scholarly publishing endeavour, and the consequent lack of any baseline against which to make comparisons.

A recent report by Larivière and Sugimoto (2018) showed a link between the monitoring of policy and its effectiveness, describing strong performance by articles supported by funders that had implemented monitoring and compliance checks for their policies. By comparison open access for works funded by Canadian funders, which did not monitor compliance, were shown to lag substantially even when disciplinary effects were taken into account.

There is also a need for critical and inclusive evaluation of open access performance that can address regional and political differences. For example, the SciELO project has successfully implemented an electronic publishing model for journals resulting in a surge of journal-mediated open access (Packer, 2009; Wang et al., 2018). Recent work by Iyandemye and Thomas (2019) showed that, for biomedical research, there was a greater level of open access for articles published from countries with a lower GDP, particularly for those in sub-Saharan Africa. This provides evidence of national or regional effects on publication cultures that lead to open access. Meanwhile, Siler et al. (2018) showed that, for the field of Global Health, lower-ranked institutions are more likely to publish in closed outlets. They suggest this is due to the cost of article processing charges showing the importance of considering institutional context when examining open access performance.

### 1.1 Change at the institutional level

We have argued (Montgomery et al., 2018) that the key to understanding and guiding the cultural changes that underpin a transition to openness is analysis at the level of research institutions. While funders, national governments, and research communities create the environments in which researchers operate, it is within their professional spaces that choices around communication, and their links to career progression and job security are strongest. Analysis of how external policy leads to change at the level of universities is critical. However, providing accurate and reliable data on open access at the university level is a challenge.

The most comprehensive work on open access at the university level currently available is that included in the CWTS Leiden Ranking (Robinson-Garsia et al., 2019). This utilises an internal Web of Science database and data from Unpaywall^2^ to provide estimates of open access over a range of timeframes. These data have highlighted the broad effects of funder policies (notably the performance of UK universities in response to national policies) while also providing standout examples from regions that are less expected (for instance Bilkent University in Turkey).

A concern in any university evaluation is the existing disciplinary bias in large bibliographic sources used to support rankings. For example, the coverages of Web of Science and Scopus were shown to be biased toward the sciences and the English language. We, and others have shown how sources can be biased to wards disciplines and languages (Mongeon and Paul-Hus, 2016) and how evaluation frameworks based on single sources of output data can provide misleading results (Huang et al., 2020a). In a companion white paper to this article we provide more details of these issues with a sensitivity analysis of the data presented here (Huang et al., 2020b). If we are to make valid comparisons of universities across countries, regions and funders to examine the effectiveness of open access policy implementation there is a critical need for evaluation frameworks that provide fair, inclusive, and relevant measurement of open access performance.

### 1.2 Challenges in evaluating institutions

Building a robust open access evaluation framework at the institutional level comes with a number of challenges. Alongside coverage of data sources are issues of scope (which institutions, what set of objects), metrics (numbers or proportions) and data completeness. Our pragmatic assessment is that any evaluation framework should be tied to explicit policy goals and be shaped to deliver that. Following from our work on open knowledge institutions (Montgomery et al., 2018) our goals in conducting an evaluation exercise and developing the framework are as follows:

1. Maximising the amount of research content that is accessible to the widest range of users, in the first instance focusing on existing formal research content for which metadata quality is sufficiently high to enable analysis
2. Developing an evaluation framework that drives an elevation of open access and open science issues to a strategic issue for all research-intensive universities
3. Developing a framework that is sensitive to and can support universities taking a diversity of approaches and routes towards delivering on those goals

In terms of a pragmatic approach to delivering on these we therefore intend to:

1. Focus on research-intensive institutions, using existing rankings as a sample set
2. Seek to maximise the set of objects which we can collect and track while connecting them to institutions (i.e., favour recall over precision)
3. Focus on proportions of open access as a performance indicator rather than absolute numbers
4. Publicly report on the details of performance for high performing institutions (and provide strategic data on request to others)
5. Report on the diversity of paths being taken to deliver overall access by a diverse group of universities
6. Develop methodology that is capable of identifying which policy interventions have made a difference to outcome measures and any ‘signature’ of those effects

## 2 Results

### 2.1 A reproducible workflow to evaluate institutional open access performance

We developed a reproducible workflow capable of quantifying a wide range of open access characteristics at the institutional level. The overall workflow is summarised diagrammatically in Figure 1. This includes a mapping of open access definitions and the Unpaywall information we used to construct them. Briefly, we gather output metadata from searches in Microsoft Academic^3^ (Sinha et al., 2015; Wang et al., 2019), Web of Science and Scopus, for each university. From this full set we gather the corresponding Crossref DOIs from the metadata of each output focusing on this set. Unpaywall is consulted to determine open access status. Detailed discussions of the data sources, precise data snapshots used, open access definitions, and technical details of the data infrastructure can be found in the Supplementary Methodology.

**Figure 1:**
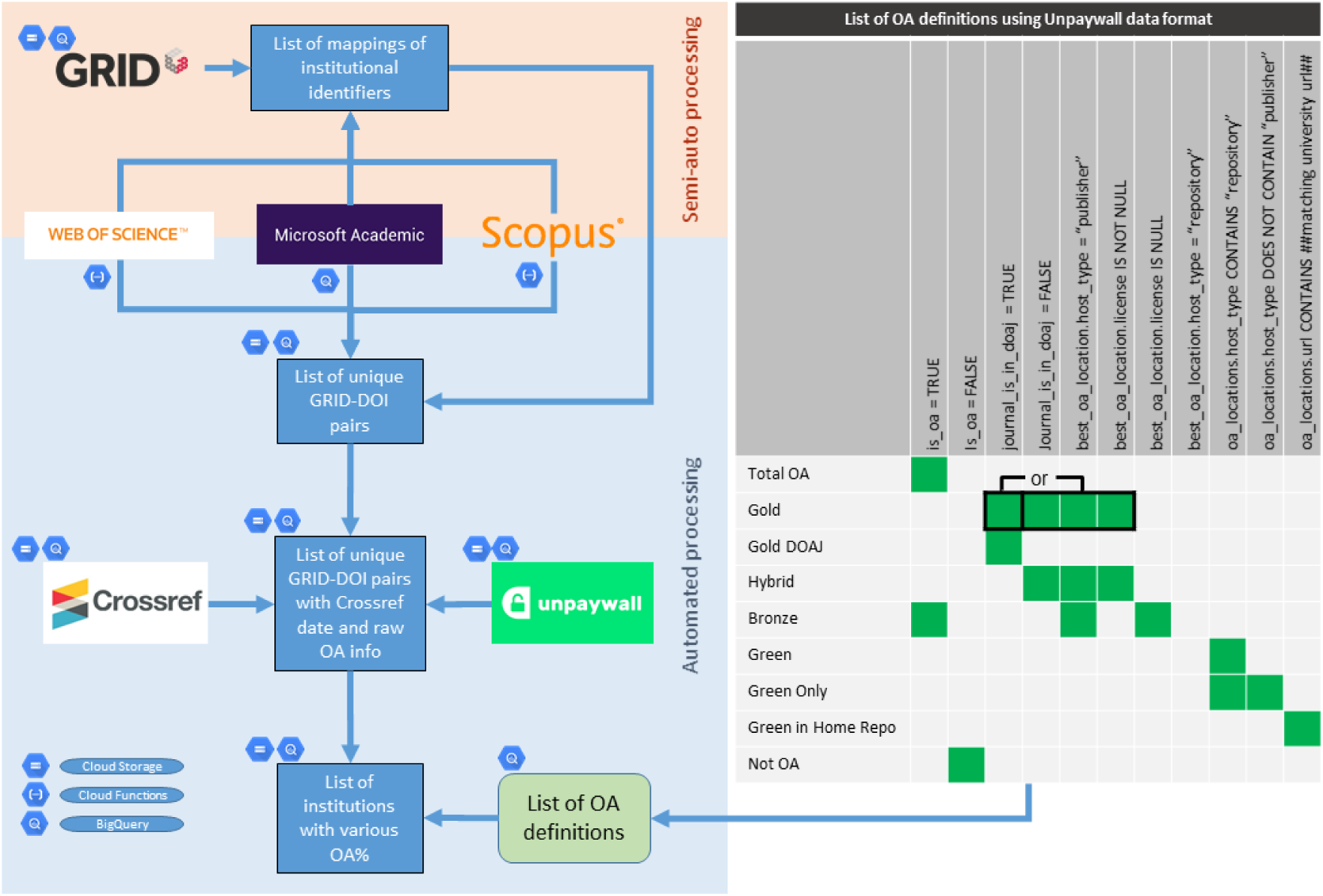
Workflow of data collection and mapping of open access definitions to Unpaywall metadata.

As we have noted previously (Huang et al., 2020a) there is a sensitivity associated to the choices in bibliographic data sources when they are used to create a ranking. For this analysis we therefore chose to combine all three datasets (i.e., Microsoft Academic, Web of Science and Scopus). In the companion white paper (Huang et al., 2020b) we provide a comprehensive sensitivity analysis on the use of these different datasets, the use of different versions of Unpaywall, and the relations between confidence levels and sample size.

Briefly, it is our view that to provide a robust assessment of open access performance the following criteria must be met:

1. The set of outputs included in each category (here institutions) and a traceable description of how they were collected must be transparently described. Provided here by a description of the data sources and the procedures used to collect DOIs for each institution (see Supplementary Methodology).
2. A clearly defined, open and auditable data source on open access status. Provided here by a defined and identified Unpaywall snapshot (see Supplementary Methodology).
3. A clearly defined and implementable description of how open access status data is interpreted. Provided here in Figure 1 and in Supplementary Methodology in the form of the SQL query used to establish open access status categories for each DOI.
4. Provision of derived data and analysis in auditable form. Provided here the derived data as open data (Huang et al., 2020c), code for the analysis of derived data as Jupyter notebooks (Huang et al., 2020d), and upstream data analysis in the form of SQL queries used (Huang et al., 2020c).

We have limited our data sharing in two ways. Firstly, we do not provide the full list of DOIs obtained from each source, due to Terms of Service restrictions. Secondly, we have not identified institutions individually except for those that fall within the top 100 globally for total open access, journal-mediated, or repository-mediated open access. The full dataset containing derived data for all institutions is available in anonymised form (Huang et al., 2020c).

### 2.2 Top 100 global universities in terms of total open access, gold open access and green open access

In Figure 2, we present the top 100 universities in our dataset for each of the categories of total open access, publisher-mediated open access (“gold”) and repository-mediated open access (“green”) for publications assigned to the year 2017 (see Supplementary Figures 1 and 2 for equivalent plots for 2016 and 2018). This is, to our knowledge, the first set of university rankings that provides a confidence interval on the quantitative variable being ranked and compensates for the multiple comparisons effect. Across this top 100 the statistical difference between universities at the 95% confidence shows that a simple numerical ranking cannot be justified. The high performance of a number of Latin American and African universities, together with a number of Indonesian universities, particularly with respect to gold open access, is striking. For Latin America this is sensitive to our use of Microsoft Academic as a data source (Huang et al., 2020b) showing the importance of an inclusive approach. The outcomes for Indonesian universities are also consistent with the latest report on country-level analysis (Van Noorden, 2019). These suggest that the narrative of Europe and the USA driving a publishing-dominated approach to open access misses a substantial part of the full global picture.

**Figure 2:**
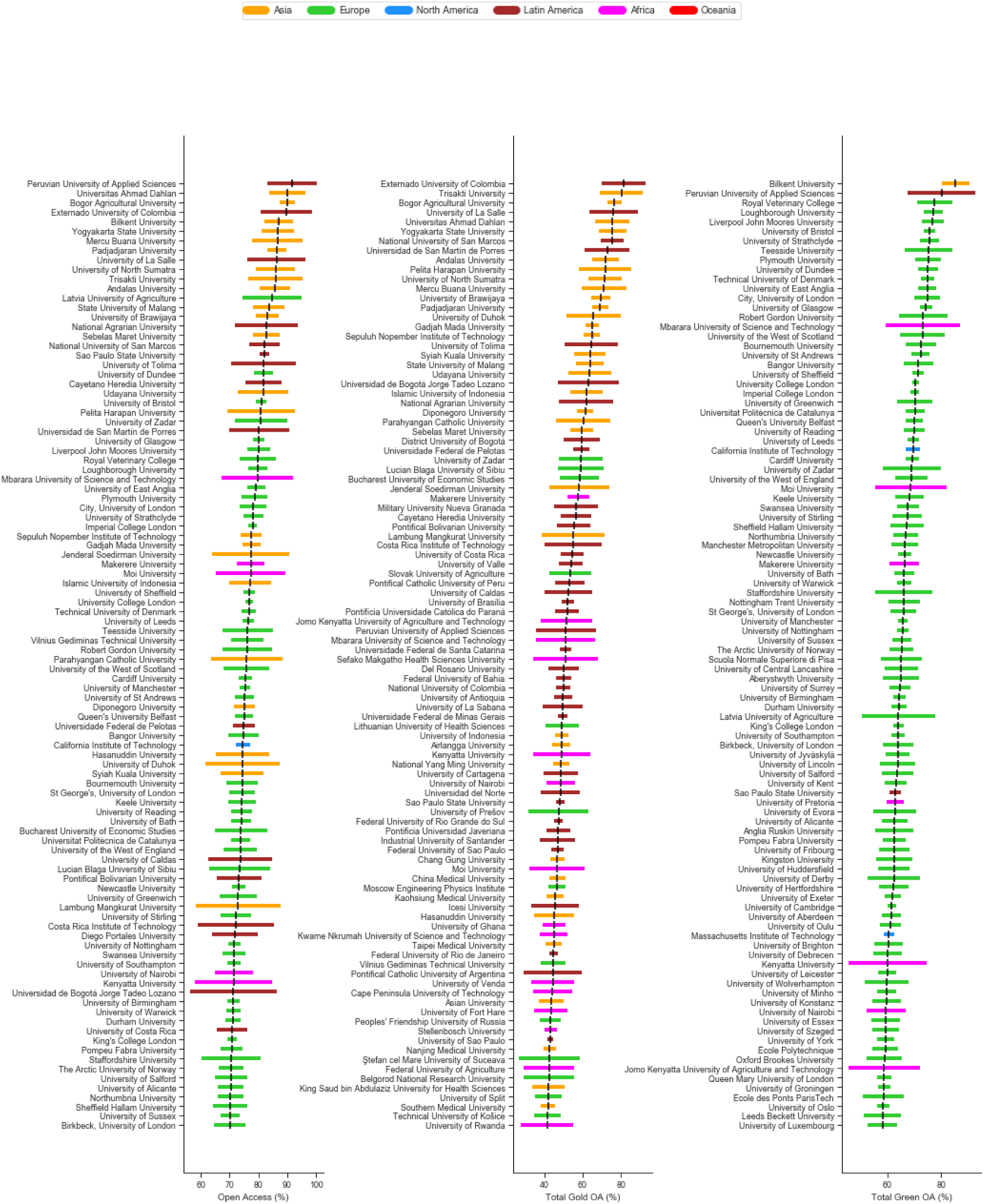
Top 100 universities in terms of performance in proportions of total open access, open access publishing (gold OA) and repository-mediated open access (green OA) for 2017.

The highest performers in terms of open access via repositories are dominated by UK universities. This is not surprising given the power of the open access mandate associated with the Research Excellence Framework to drive university behaviour. It is perhaps interesting that few US universities appear in this group (with CalTech and MIT the exceptions). This suggests that while the National Institutes of Health mandate has been very effective at driving open access to the biomedical literature limited inroads have been made into other disciplines in the US context, despite the White House memorandum. As was seen in the Leiden Ranking, Bilkent University from Turkey also emerges as a stand-out performer.

### 2.3 The global picture and its evolution

The levels of total open access, publisher-mediated open access and repository-mediated open access for 1,207 universities for publications in 2017, grouped by country, can be found in Supplementary Figure 3 (and for other years given in Supplementary Figures 4-5). Countries are ordered by the median total open access percentage. Amongst countries with a large number of universities in the dataset the UK is a clear leader with Indonesia, Brazil, Columbia, the Netherlands, and Switzerland showing a strong performance. Supplementary Figures 6-8 show results grouped by regions.

There are high performing universities in Latin America (i.e., Peru, Costa Rica, Columbia, Chile, Brazil) and Uganda as well as a range of European countries. Latin American countries owe their performance in large part to open access journals (“gold”) whereas European countries see a more significant contribution from repository based open access (“green”). Many countries have universities that are high performers in terms of the proportion of open access, while overall country performance can be linked to policy mandates and infrastructure provision.

To examine the global picture for the 1,207 universities in our dataset and to interrogate different paths to open access we plot the overall level of repository mediated (“green”) and publisher mediated (“gold”) open access for each university over time coloured by region as previously. Figure 3 presents the results for 2017 (with changes over time shown in the animated version, see also Supplementary Figure 9 for each year).

**Figure 3:**
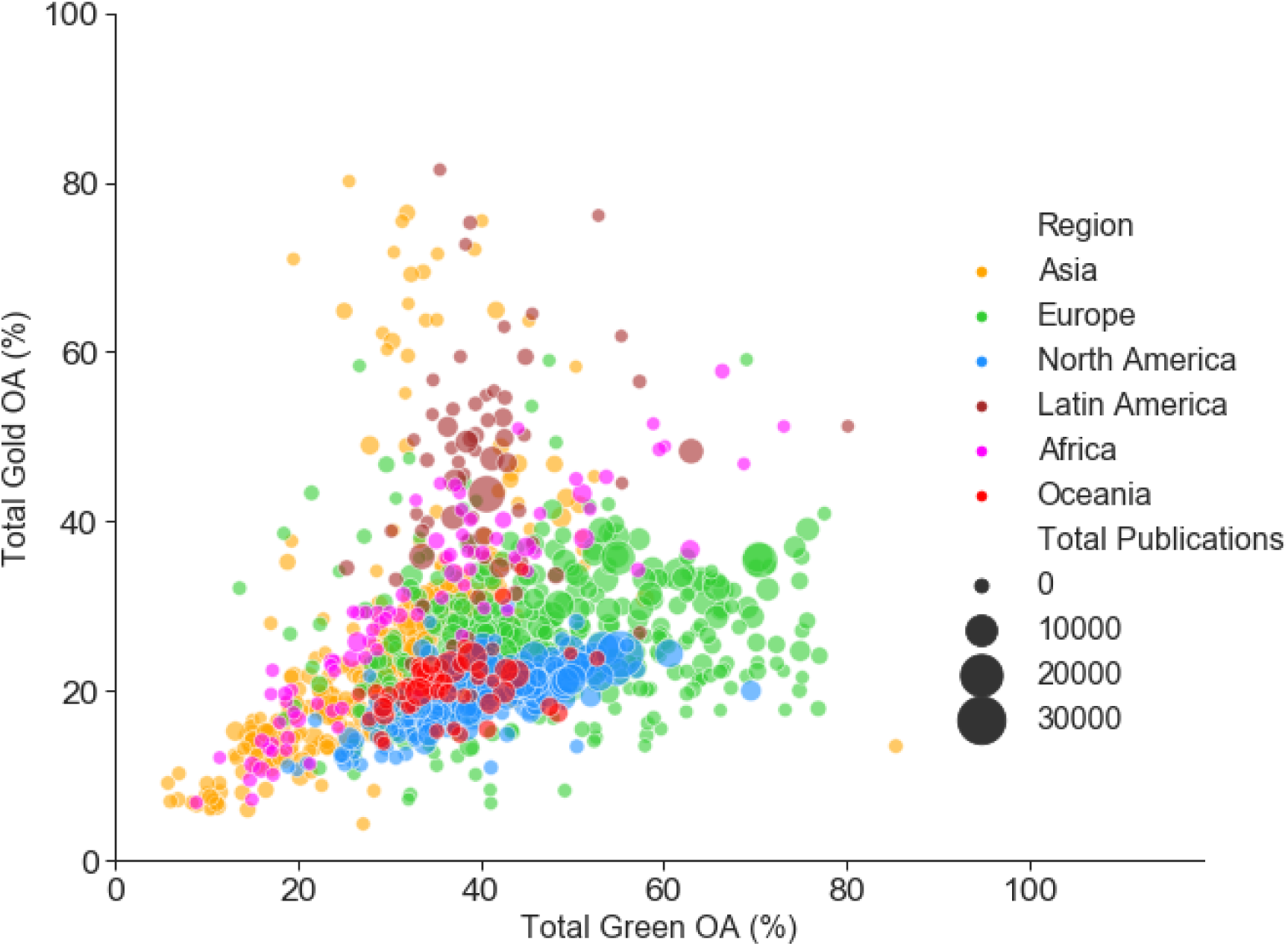
Open access publishing (gold OA) vs repository-mediated open access (green OA) by institution for 2017 (and 2007-2018 for animated version). Each point plotted is a university, with size indicating the number of outputs analysed and colour showing the region. Articles can be open access through both publishing and repository routes so x and y values do not sum to give total open access.

Overall universities in Oceania (Australia and New Zealand) and North America (Canada and the US) lag behind comparators in Europe (on repository-mediated open access) and Latin America (on open access publishing). Asian universities are highly diverse. As seen in Figure 2 there are some high performers in the top 100s, particularly for open access publishing, but many also lag behind. Africa is also highly diverse but with a skew towards high performance, with an emphasis on open access publishing. This may reflect our sampling which is skewed towards institutions with the largest (formally recorded) publishing volumes, many of which receive significant portions of their funding from international donors with strong open access requirements. Latin American institutions show high levels of open access publishing throughout the period illustrated. This is due to substantial infrastructure investments in systems like SciELO starting in the 1990s.

### 2.4 The effects of policy interventions

If our goal is to provide data on the effectiveness of interventions then our analysis should be capable of identifying the effects of policy change. In 2012 the UK Research Councils, following the Finch Report, provided additional funding to individual universities to support open access publishing. The amount of additional funding relates to existing research council funding. In Figure 4a we show the annual change in open access publishing for three UK universities with the largest additional funding and three with significantly less additional funding (Lawson, 2018). In either 2012 or 2013 we see a jump in open access publishing across all the universities. This effect is most clear for the hybrid open access contribution to gold (Supplementary Figures 10a and 10b). As the additional funding tails off in 2015 the rate of growth falls back.

**Figure 4:**
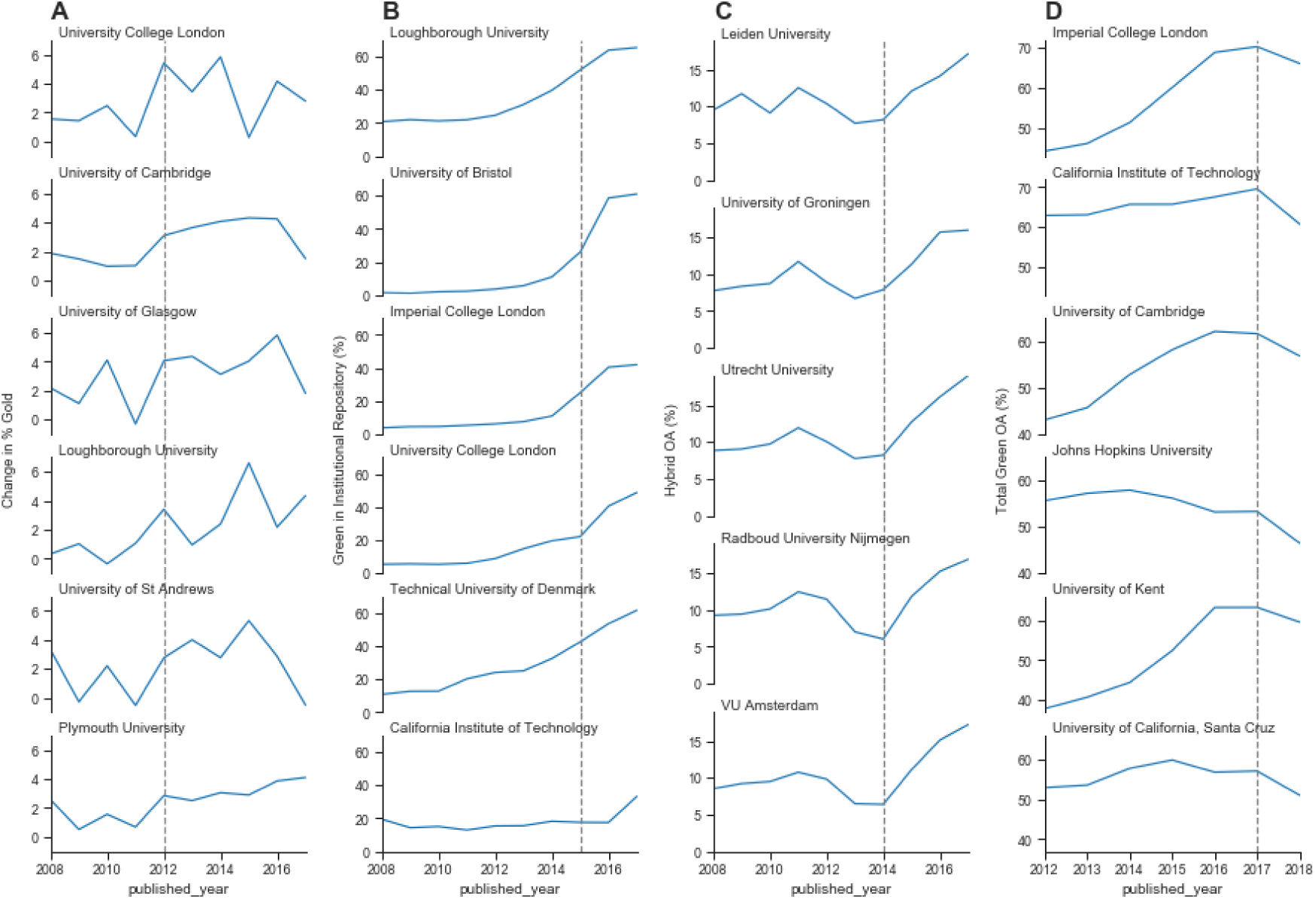
Monitoring the effect of policy interventions for selected groups of universities. Panel A shows the annual change in percentage (rolling current year percentage minus the previous year percentage) of gold OA for six UK universities. The first three universities are those with larger additional funding in contrast to the last three universities who received less additional funding. Panel B shows the annual percentage of green OA through the home institutional repositories of four UK universities compared to high performing universities from elsewhere. Panel C shows the annual percentages of hybrid OA at six universities in the Netherlands. Panel D shows three pairs of UK and US universities, selected based on having a similar size and level of green OA. The annual percentages of total green OA are depicted for each university.

Figure 4b shows the growth of content in UK university repositories from 2000-2017, compared to two universities from other regions. In 2015 deposit of a research output in a repository became a requirement for eligibility for including in the UK Research Excellence Framework. This policy shift was profound because it relates to an assessment exercise and funding which covers all disciplinary areas and all universities. The dominance of the top 100 for both overall open access and repository-mediated open access by UK universities as well as the approach to 100% coverage being made by such a large number of universities is driven in large part by that intervention.

Figure 4c focuses on the take-up of hybrid open access publishing options in the Netherlands following deals with Springer in 2014, and Wiley in 2016. The consistent dip in hybrid adoption in the Netherlands to 2014 does not have an obvious explanation except perhaps that researchers were waiting to see the result of negotiations. Across the Netherlands levels of publishing in hybrid open access journals show a sharp turn of increase from 2014 onwards with a less pronounced effects (more smooth increases) for publishing in pure open access (see Supplementary Figures 10c and 10d).

Finally, in Figure 4d we show the effect of subtle differences in policy relating to acceptable embargo periods. UK Research and Funding Council polices have been aggressive in reducing embargo lengths mandating six months for STEM subjects and twelve months for HSS subjects. The effect of embargoes can be seen in data for repository mediated open access as a dip in the most recent years of publication. Using Unpaywall data from late 2019 we see a dip in repository-mediated open access performance for UK universities in 2018 but a limited effect on 2017. By comparison with three of the highest performing US universities, comparable in size and overall ranking (see Figure 2), we see an extended dip in performance, indicative of an acceptance of longer embargoes.

### 2.5 Different institutional paths towards open access

In Figures 3 and 4 we see evidence of different paths towards open access, depending on the context and resources. The idea of mapping these paths is shown explicitly for a subset of universities in Figure 5. This shows the paths taken by two sets of UK universities and a selection of Latin American institutions over time. For UK universities that received substantial funding from the UK research councils for open access publishing three examples are shown. An alternate route, emphasising repository-mediated open access, is seen for three universities that received less additional funding.

**Figure 5:**
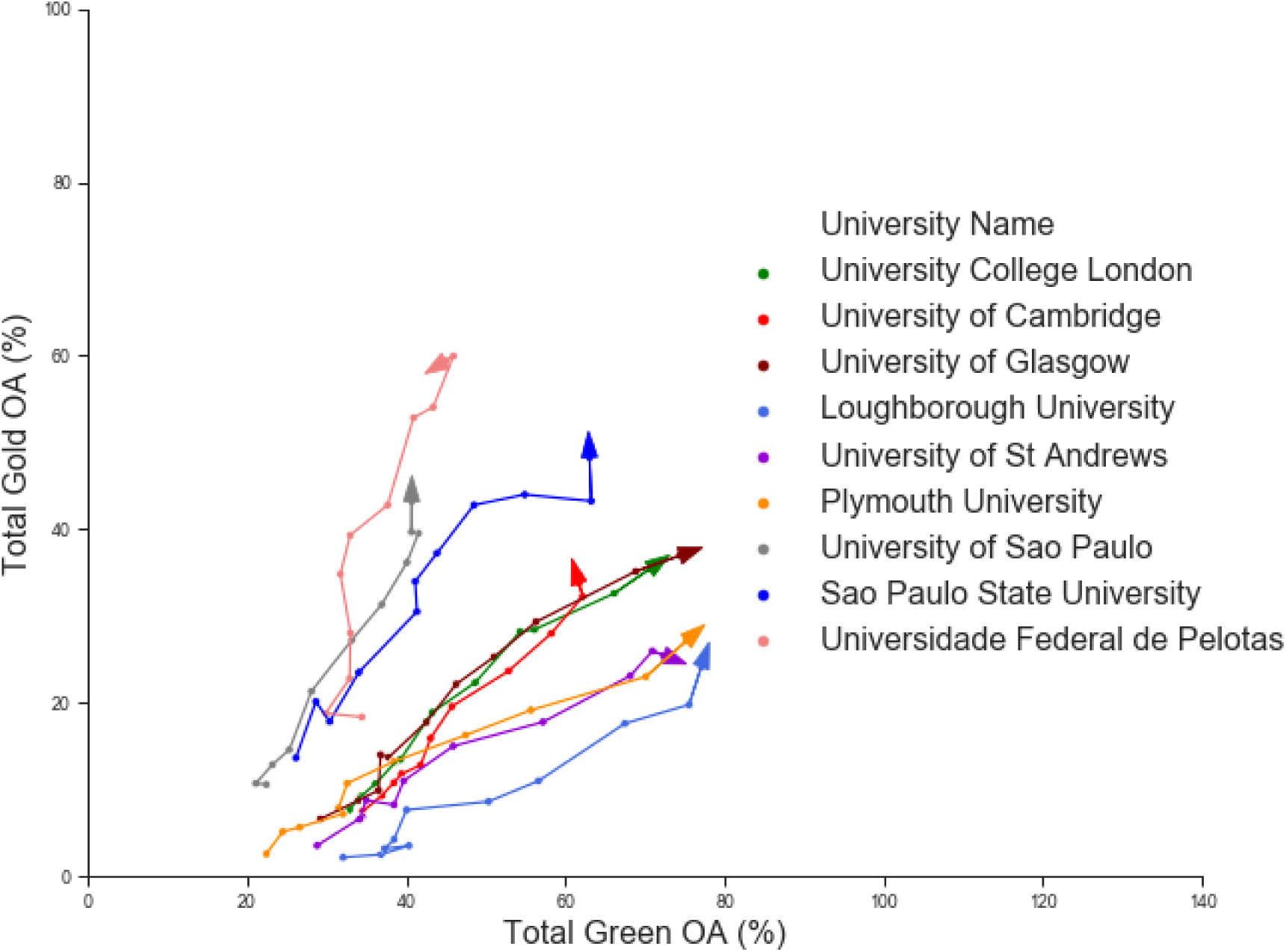
Comparing different paths to open access (gold OA versus green OA) for a selected set of universities.

In contrast the Latin American institutions already have high levels of open access publishing at our earliest time point, as discussed earlier. However, our data suggests a fall in overall open access amongst Latin American universities from 2012 onwards which we ascribe to an increased pressure to publish in “international” journals which are often subscription based, and for which Latin American scholars are reluctant or unable to pay hybrid Author Processing Charges.

## 3 Discussion

### 3.1 Implications for evaluating open access and limitations

Previous work has been mostly limited to one off evaluations and provided a limited basis for longitudinal analysis. Our analysis process includes automated approaches for collecting the outputs related to specific universities, and the analysis of those outputs. Currently the addition of new universities, and the updating of large data sources is partly manual but we also expect to automate this in the near future. Along with the nature of this article this will provide an updatable report and longitudinal dataset that can provide a consistent and growing evidence source for open access policy and implementation analysis.

While it is clear (Huang et al., 2020b) that our analysis has limitations in its capacity to provide comparable estimates of open access status across all universities, our approach does provide a reproducible and transparent view of overall global performance. There are challenges to be addressed with respect to small universities and research organisations and we have taken a necessarily subjective view of which institutions to include (see Supplementary Methodology). Our approach systematically leaves out most universities with very small number of outputs (i.e., less than 100 outputs), and universities with very extreme open access proportions as these are the universities for which we have less statistical confidence in the results. This is also in-line with our intended focus on research-intensive universities. These small institutions are of significant interest but will require a different analysis approach. We include the full set of institutions in our data set in Supplementary Figures 3 to 8.

We have used multiple sources of bibliographic information with the goal of gaining a more inclusive view of research outputs. Despite this there are still limitations in the coverage of these data sources, and a likely bias towards STEM disciplines. In addition the focus of Unpaywall on analysis of outputs with Crossref DOIs means that we are missing outputs for disciplines (humanities) and output types (books) where the use of DOIs is lower. In addition, due to the nature of this work and to limitations on the use of Web of Science and Scopus APIs, we have collected data from these two sources over a period of time. Although we expect such changes to be small those effects are not clearly represented in our data. For other data sources we are able to precisely define the data dump used for our analysis, supporting reproducibility as well as modified analyses.

### 3.2 Requirements and approaches for improving open access evaluation

There have been many differing assessments of open access performance over the past 10-15 years. Many of the differences between these have been driven by details in the approach. This combined with limited attention to reproducibility has led to confusion and a lack of clarity on the rate and degree of progress to open access (Green, 2019). As noted above we believe that a minimum standard should be set in providing assessments of open access to support evidence based policy making and implementation (see Section 2.1).

With such a minimum standard in hand we can clearly identify areas to improve open access performance assessments. There is significant opportunity for improving the data sources on sets of outputs and how they can be grouped (e.g. by people, discipline, organisation, country, etc). Improvements to institutional identifier systems such as the Research Organisation Registry, increased completeness of metadata records, particularly that provided by publishers via Crossref on affiliation, ORCIDs and funders, and enhancing the coverage of open access status data (for instance by incorporating data from CORE and BASE), will all enhance coverage. There are also opportunities to expand the coverage by incorporating a wider range of bibliographic data sources.

Broadly speaking we would advocate for the evidence base for open access policy and implementation to be built on open and transparent data. While the majority of sources we have used in this report are open (Microsoft Academic, Crossref, Unpaywall) we have elected to supplement this with data from two proprietary sources, Web of Science and Scopus. Our argument for taking this approach has been laid out separately (Huang et al., 2020a; 2020b). Here we note that our sensitivity analysis (Huang et al., 2020a; 2020b) shows that where the goal is to describe sector-wide trends and movements (as opposed to individual university performance), that the difference between using the open Microsoft Academic data alone and incorporating proprietary data is modest.

### 3.3 Implications for policy intervention and implementation

Our results have significant implications for the details of policy interventions. Firstly, we have demonstrated the ability to detect signals of policy interventions in the behaviour of institutions. We see clear effects and results arising from the efforts of national funders and policy makers, particularly in the United Kingdom. The combined policy change and funding provided by the UK Research Councils in 2012 is associated with a increase in the level of open access publishing, and the level of increase appears to be associated with the level of funding provided. Similarly the requirement for outputs to be deposited in a repository for eligibility for the 2021 Research Excellence Framework is associated with substantial increase in repository-mediated open access around 2015.

Our results also may have implications for deciding on the effectiveness of directly funding open access publishing. It is perhaps surprising to some readers that the overall levels of open access publishing in the UK are not higher. Specific funders, most notably the Wellcome Trust, have achieved very high levels of open access for articles from research they support through the provision of funding for open access publication. In addition the UK Research Councils invested significant resources in supporting gold open access. However, these have not translated to high levels of open access publishing across the full diversity of outputs of UK institutions. The majority gains over the past five years have come from repository-mediated open access.

In the animated version of Figure 3 (or the version over several years in Supplementary Figures) there is a clear signal of saturation with respect to open access publishing (gold open access) for European and North American universities. With few exceptions, institutions do not achieve levels of gold open access greater than 40% and this level is stable from 2014-2018. Similarly in Figure 4 we see evidence of shifts in response to stimuli (funding and policy interventions) which then stabilise. Even those UK universities with very high levels of repository open access (green) see a slowing down of the rise in levels a few years after the Research Excellence Framework policy intervention.

These signals suggest that the last few percent may be very difficult, and possibly expensive to achieve. There will always be areas and cases where open access is challenging. Achieving “100%” may require a tighter definition of what should be in scope. For those areas where we see signals of saturation much lower than 100% these are likely signals of the complexity of the system, and of large categories of outputs where open access is harder to achieve, or the motivation of institutions (including authors, libraries, and other support staff) to achieve it is lower.

Alongside this, the continued leadership of Latin American institutions on open access publishing levels is the continuation of a trend set more than a decade ago through the provision of publishing infrastructures. Taken alongside the clear response in the Netherlands for hybrid open access in response to publish and read agreements this suggests that increasing levels of open access publishing through article processing charges is potentially expensive compared to the costs of providing infrastructure.

Another interesting natural experiment is how the strength of funder actions is associated with overall change in levels of open access. In the Netherlands and the UK in particular, but also in the US, where funder policies have moved from encouragement, to mandates, to monitoring with sanctions for non-compliance there are substantial shifts in overall levels of open access. By contrast, in countries where policy remains effectively at the level of a recommendation, such as Australia, levels of open access lag significantly. Recent increases in reporting requirements by Australian funders might therefore be expected to lead to a detectable signal over the next 12-24 months.

Perhaps most interestingly in light of the debates surrounding Plan S is the evidence that it is universities in Latin America and Africa where levels of open access publishing are at their highest. As noted above, in Latin America this is a strong signal of the effectiveness of infrastructures such as SciELO in supporting uptake of open access practices. In the case of Africa there may be effects of funder requirements (with funders such as the Bill and Melinda Gates Foundation and Wellcome Trust that have strong open access requirements playing a significant role) as well as disciplinary spread. In both cases we are likely to have a limited view of the full diversity of research outputs due to their poor capture in information systems from the North Atlantic.

## 4 Conclusion

The evidence-base for policy development and implementation for open access has been hampered by a lack of consistency in analysis results and clarity on how those results were obtained. In particular it has been challenging to provide longitudinal and transparent results to monitor the effects of policy and support interventions. While not all readers will agree with the choices we have made in implementing an analysis process we aimed to provide sufficient transparency and reproducibility to allow for both replication, critique and alternative approaches to this analysis. This can underpin a higher quality of debate and policy development globally, and aid in learning from successes in other regions.

Our analysis of open access performance by research-intensive universities highlights the importance of robust policy and support in driving change. Geographies where there is a long history of infrastructure provision, such as Latin America, show very high levels of open access publishing. The United Kingdom is a particular case where there is consistently very high levels of open access, particularly that provided through repositories, in response to a strong and well supported policy environment. We also see that different institutions may choose to take different paths to delivering open access depending on resources, culture and systems in place.

The value of analysis at the level of universities is that we gain a picture of open access performance across a diverse research ecosystem. We see differences across countries and regions, and differences between universities within countries. Overall we see that there are multiple different paths towards improving access, and that different paths may be more or less appropriate in different contexts. Most importantly, while further research is needed to unpick the details of the differences in open access provision, we hope this work provides a framework for enabling that longitudinal analysis to be taken forward and used wherever it is needed.

## Supporting information

Animated version of Figure 3

Animated version of Figure 5

## 5 Acknowledgements

This work was funded by the Research Office of Curtin University through a strategic grant, the Curtin University Faculty of Humanities, and the School of Media, Creative Arts and Social Inquiry.

## Supplementary Methodology

### 1 Technical infrastructure, reproducibility and provenance

Our technical infrastructure is constructed based on the aim to make openly available both the data and analysis code as much as possible. The data infrastructure is currently based in Google Cloud Platform, mainly utilising services in Google Storage, Google Functions, and Google Bigquery. Google Functions are used to extract data from the data sources’ APIs. The raw data is then stored on Google Storage for further processing. Google Bigquery is then used to merge and manipulate the raw data to process derived data in formats we require for further analysis.

Derived data used for analysis can be found at Zenodo (Huang et al., 2020b). Updated datasets will also be provided and will be found at the same location. Raw data is not provided to preserve the anonymity of institutions and respect the terms of service of data providers. The SQL queries and code used to generate the derived datasets are described below and available via Zenodo.

The main article, Supplementary Figures and this Supplementary Methodology were prepared as Jupyter notebooks to provide all analysis and visualisation code and maximise reproducibility. These notebooks are available at Github and Zenodo (Huang et al., 2020c). All manipulation of derived data after import is explicitly conducted in the notebook. The notebook utilises a library for generating visualisations, which is provided with the notebooks. The only data manipulations performed by the visualisation library are to filter and re-shape the data for graphing.

Where possible we use a publicly available or defined data dump for our sources. In this article data from Crossref, Unpaywall, GRID and Microsoft Academic were available as data dumps. The dumps used in this version (submitted version 19 March 2020) of these articles are as follows:

| Data Source | Dump Identity | Availability | License |
| --- | --- | --- | --- |
| Crossref | December 2019 | CR Metadata Plus Members<br>https://www.crossref.org/help/metadata-snapshots/ | cc0 |
| Unpaywall | 22 November 2019 | By request<br>http://unpaywall.org/products/snapshot | cc0 |
| GRID | Release 2019-12-10 | http://doi.org/10.6084/m9.figshare.11353022 | cc0 |
| Microsoft Academic Graph | 3 October 2019 | Via Azure<br>https://docs.microsoft.com/en-us/academic-services/graph/get-started-setup-provisioning | ODC-By |

### 2 Data sources

We integrate a variety of data sources in our data workflow to generate open access scores for a large set of universities. These sources include the following:

#### 2.1 Web of Science

Web of Science is a large pay-walled online scientific citation indexing service, maintained by Clarivate Analytics, with the coverage of more than 90 million records and 1.4 billion cited references. It is often harvested by universities to build-up their internal research information database. And, with much criticism, it is also used by various university rankings to evaluate performance (e.g., Academic Ranking of World Universities, CWTS Leiden Ranking, U-Multirank, etc). It stands as an important tool for various stakeholders of the academia. Web of Science includes a number of databases with varying levels of accessibility and information. For this study, we utilise the “organization-enhanced” search functionality to extract the list of publication metadata (hence, the corresponding DOIs) for each institution of interest (via our local access) from the Web of Science Core databases. Our access to Web of Science Core is restricted by our institutional subscription contract, which provides access to the following:

- Science Citation Index Expanded (SCI-EXPANDED) –1972-present
- Social Sciences Citation Index (SSCI) –1972-present
- Arts & Humanities Citation Index (A&HCI) –1975-present
- Conference Proceedings Citation Index-Science (CPCI-S) –1990-present
- Conference Proceedings Citation Index-Social Science & Humanities (CPCI-SSH) –1990-present
- Book Citation Index– Science (BKCI-S) –2005-2012
- Book Citation Index– Social Sciences & Humanities (BKCI-SSH) –2005-2012
- Emerging Sources Citation Index (ESCI) –2015-present
- Current Chemical Reactions (CCR-EXPANDED) –1985-present
- (Includes Institut National de la Propriete Industrielle structure data back to 1840)
- Index Chemicus (IC) –1993-present

#### 2.2 Scopus

Scopus is an abstract and citation database launched by Elsevier in 2004. It provides subscription access and also produces a range of quality measures such as h-index, CiteScore and SCImago Journal Rank. For the purpose of our current work, we match each institution to its Scopus Affiliation ID and, subsequently, access the metadata of all publications related to each institution (again, via local access). The DOI, if existent, is extracted from each publication’s metadata.

#### 2.3 Microsoft Academic

Microsoft Academic, re-launched in 2016, is a replacement of the phased-out Microsoft Academic Search. It is a free public search engine for the academic literature and uses the semantic search technology developed by Microsoft Research. The database provides Affiliation Entity IDs for institutions. We utilise a snapshot of Microsoft Academic database to extract publication metadata related to each institution.

#### 2.4 Times Higher Education World University Rankings

This is an annual ranking produced by the Times Higher Education (THE) magazine. It is one of the most followed university rankings, together with the Academic Ranking of World Universities and Quacquarelli Symonds (QS) World University Rankings. The major components of the THE Ranking include its reputation survey and the citations data from Web of Science. As a mean for comparisons, we selected the Top 1000 universities in the 2019 THE Ranking as our primary sample for calculating OA scores. This is supplemented with additional universities for countries with limited coverage in the primary sample. Subsets of universities are selected for longitudinal studies.

#### 2.5 Unpaywall

Unpaywall is a browser extension for finding free legal versions of paywalled research publications. It currently covers more than twenty-two million free scholarly articles and provides a large number of metadata related to OA, such as journal OA status (via DOAJ) and open license information. It has recently been integrated into the Web of Science and Scopus databases. For this study, each DOI of interest is matched with its metadata in Unpaywall for calculating the various OA status. Snapshots of the Unpaywall database are collected as part of data processing of this project.

#### 2.6 Crossref

Crossref is a not-for-profit official DOI registration agency of the International DOI Foundation. It is the largest (in terms of number of DOIs assigned) DOI registration agency in the world. It also provide JSON structured information surrounding each of its DOI, such as various related issue dates, and links between distributed content hosted at other sites. Our data collection process have resulted in several sanpshots of the Crossref database. We primarily view Crossref DOIs as the basis for coverage of all global outputs in our study. We use the `issued.date` element in Crossref as the standardised indicator for publication year for each output in our data.

#### 2.7 Global Research Identifier Database (GRID)

This is an open access database of educational and research institutions worldwide. It assigns a unique GRID ID to each institution and, where applicable, to each level of the institutional hierarchy. The metadata includes information such as geo-coordinates, websites and name variants. These identifiers are adapted in our study to link the various bibliographic data sources and to unify the identification system.

### 3 Description of data workflow and selection criteria

As discussed in the main article, our pragmatic approach is to include the widest coverage of outputs for each of the universities under consideration. This implies defining a *target population* for all potential research outputs, which is no trivial task. For this study, we choose to consider the set of all research outputs with Crossref DOIs as this target population. This is identified as the most practical approach that allows tracking and disambiguation of research objects using persistent identifiers. At the same time, it provides processes for both the standardisation of publication dates and the use of Unpaywall’s OA information.

We use universities listed in the top 1000 of the Times Higher Education World University Rankings as an initial sample for which to collect data. We then supplemented this with additional institutions, focussing on the United Kingdom and the United States. Finally we added additional universities in countries where our original sample had one or two universities. For each of these countries we added a small number of additional universities, prioritising those with the largest number of research outputs recorded in Microsoft Academic.

Given that Microsoft Academic, Web of Science and Scopus have different internal institutional identifier systems, the next step is to map these identifiers. We first map each university to its unique ID in the Global Research Identifier Database (GRID). Subsequently, these universities’ internal identifiers for Microsoft Academic, Web of Science and Scopus are matched against the corresponding GRID IDs. This is trivial in the case of a Microsoft Academic database snapshot as each institution in its database is already matched against the corresponding GRID ID. For Web of Science and Scopus, manual website searches are required to retrieve Web of Science Organisation-Enhanced names and Scopus Affiliation IDs, respectively. ***Universities not identifiable in at least one of the three bibliographic data sources are not processed further***.

Queries were run via the respective APIs against Web of Science (via Organisation-Enhance name search) and Scopus (via Affiliation ID search) to extract metadata of all outputs affiliated to each university for the time frames 2000 to 2018. These are matched against outputs from a Microsoft Academic snapshot to result in a comprehensive set of outputs for each university. Subsequently, these are filtered down to include only objects with Crossref DOIs. This current set of universities is then further expanded to include additional universities from countries that had low representations in the initial sample, and goes through the same data collection process.

All collected Crossref DOIs are matched against an Unpaywall database snapshot for their open access information. This allows us to calculate total numbers for various modes of open access (e.g., number of Gold OA publications) for each university across different timeframes (using the “year” component of the Crossref “issued date” field). The Unpaywall information used to determine various open access modes is as displayed in Figure 1 of the main text. Crossref DOIs not found in Unpaywall are defaulted to be not open access and Crossref DOIs that do not have an “issued date” are removed from the process.

A comprehensive sensitivity analysis on the use of different sources for gathering research outputs, use of different Unpaywall versions, and the relations between confidence levels and sample size is provided in the companion white paper Huang et al. (2020b). There are changes based on which specific Unpaywall snapshot is used. This is partly due to real changes (e.g. release of works from repositories after embargo) and due to changes within the Unpaywall data system (examples include changes in upstream data sources such as journal inclusion or exclusion in the Directory of Open Access Journals (DOAJ), and internal changes such as improved repository calling or wider journal coverage). As this is a product of gradually improving systems underpinning Unpaywall we use the most recent available snapshot to provide the most up to date data in a reproducible and identifiable form in the main article.

To make comparable and fair analysis across universities, we have taken a necessarily subjective view on which universities to include in Figures 2 and 3 (along with their supplementary figures). We use the Šidák correction to control for the familywise error rate in multiple comparisons (this essentially results in individual confidence intervals and margins of error being evaluated at 99.95%). Institutions with margin of error greater than 17, for any of total open access, gold open access or green open access, were also removed from data used to generate various figures. This is in addition to the conventional conditions for normal approximation (see Huang et al., 2020b).

#### 3.1 Overview of the data workflow and main tables

Within our Google Cloud Platform environment we maintain two sets of tables. The latest snapshot of each part of the dataflow is distinguished as `_latest` and specific snapshots are named for the date of production. We maintain a set of views within the Google Cloud BigQuery platform that are setup to query the latest available data. For this article we have created a specific snapshot, and that data is shared. Future versions of the shared dataset will provide updated snapshots.

The overall flow of data and the main tables and queries are described in Figures SM1 and SM2. The key table in our workflow is the `institutions` table. Figure SM1 details the main elements of the workflow that generates the `institutions` table. Figure SM2 details the downstream processing used to generated the derived datasets used in our analyses.

**Figure SM1:**
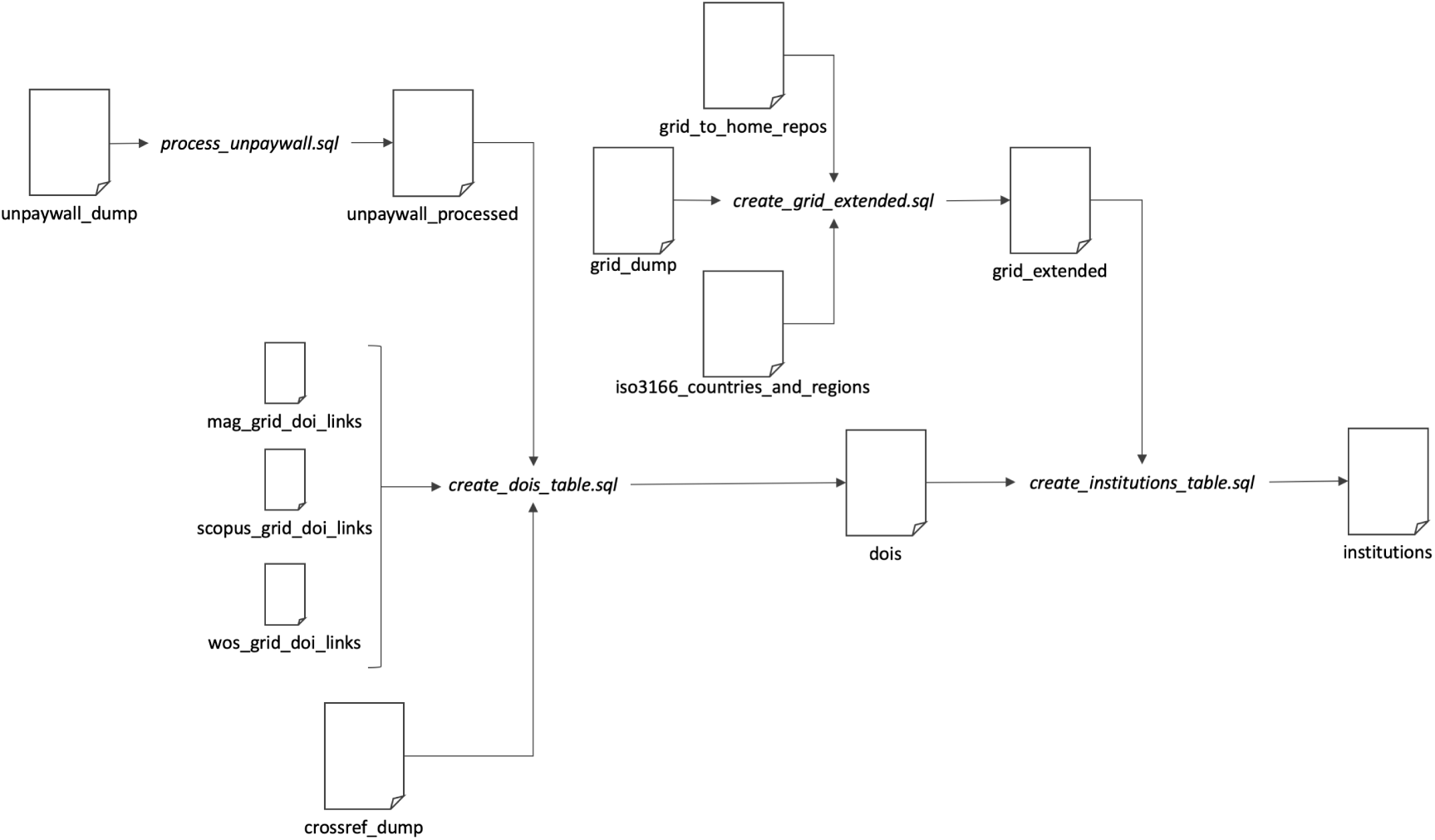
The generation of the institutions table.

**Figure SM2:**
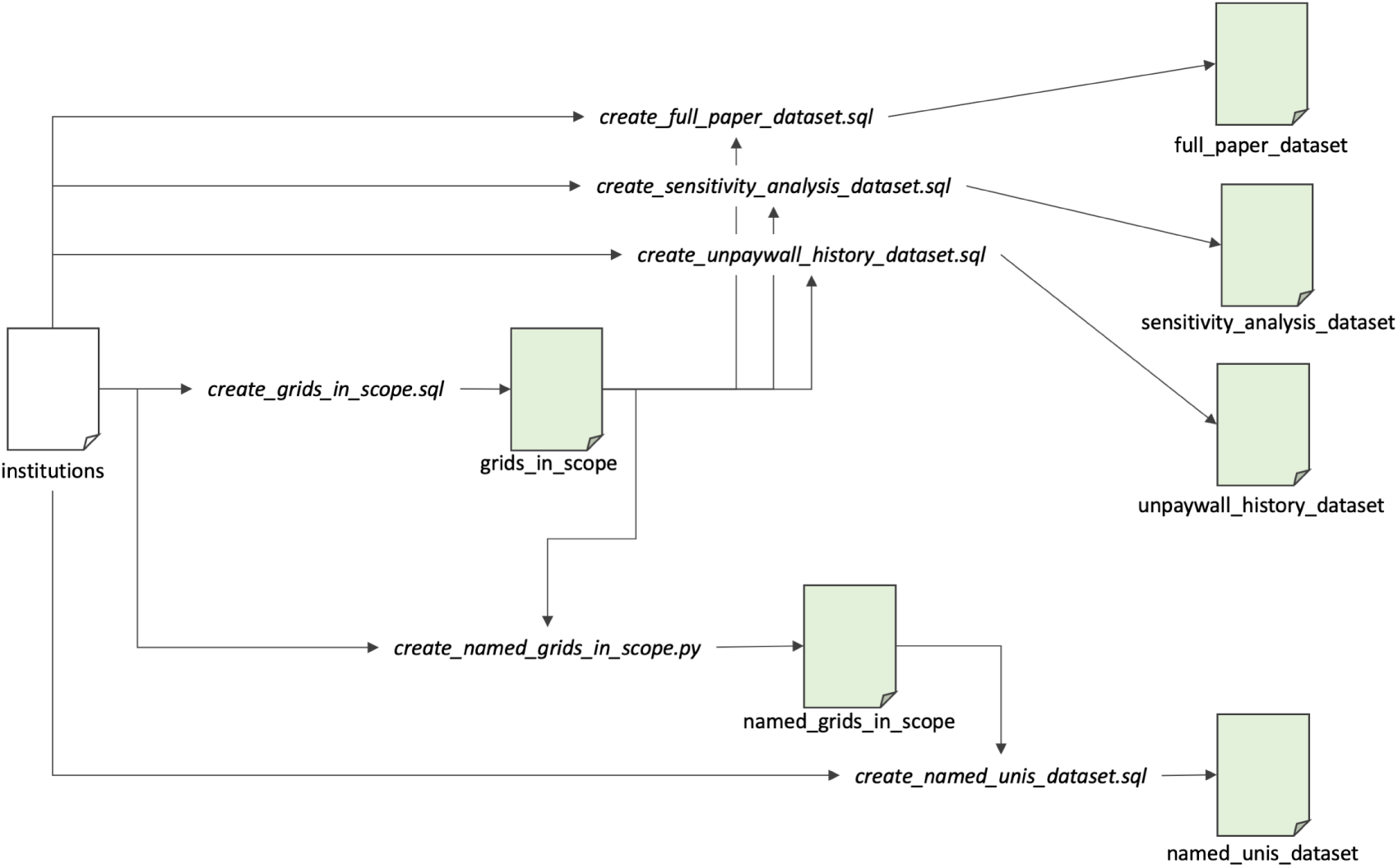
The generation of the derived datasets from the institutions table. Those datasets which are part of the publicly shared derived data are shown in green.

#### 3.2 Modes of open access

As summarised in the main article, we query each of three bibliographic data sources (Web of Science, Scopus and Microsoft Academic) for its list of research output affiliated to a given university, from years 2000 to 2018. Subsequently, this is filtered down to all objects with Crossref DOIs (by mapping against a Crossref data snapshot) and matched against Unpaywall metadata. We use the aggregated sets of DOIs for each year of publication (as per “issued date” in Crossref) to compute the counts for various OA modes using data from Unpaywall. The details of how different OA characteristics are calculated is shown in Figure 1 in the Results section of the main article. The details of the SQL query used to categorise OA status can be found below.

While there is a large body of literature on OA, the definitions of OA are quite diverse in detail. Policy makers and researchers may choose to use the OA terminology in different ways. Popular discrepancies include the coverage of journals without formal license of reuse and articles only accessible via academic social medias or illegal pirate sites. We use the following definitions for the modes of OA determined as part of our data workflow:

- **Total OA:** A research output that is free to read online, either via the publisher website or in an OA repository.
- **Gold:** A research output that is either published in a journal listed by the Directory of Open Access Journals (DOAJ), or (if journal not in DOAJ) is free to read via publisher with any license.
- **Gold DOAJ:** A research output that is published in a journal listed by DOAJ.
- **Hybrid:** A research output that is published in a journal not listed by DOAJ, but is free to read from publisher with any license.
- **Bronze:** A research output that is free to read online via publisher without a license.
- **Green:** A research output that is free to read online via an OA repository.
- **Green Only:** A research output that is free to read online via an OA repository, but is not available for free via the publisher.
- **Green in Home Repo:** A research output that is free to read online via the matched affiliation’s institional repository.

It should be noted that these definitions are not always mutually exclusive in coverage. For example, an article can be both Gold OA and Green OA. On the other hand, the set of all Gold OA and the set of all Green Only OA do not have any common element by definition. In the main text of this article we only report the categories: Total OA, Gold, Hybrid, Green, Green in Home Repo. A sensitivity analysis of the use of alternative categories of OA can be found in the companion white paper.

The full query that processes the Unpaywall data to processed open access status is as follows and can also be found in the Data and Queries package at Zenodo (Huang et al., 2020b).

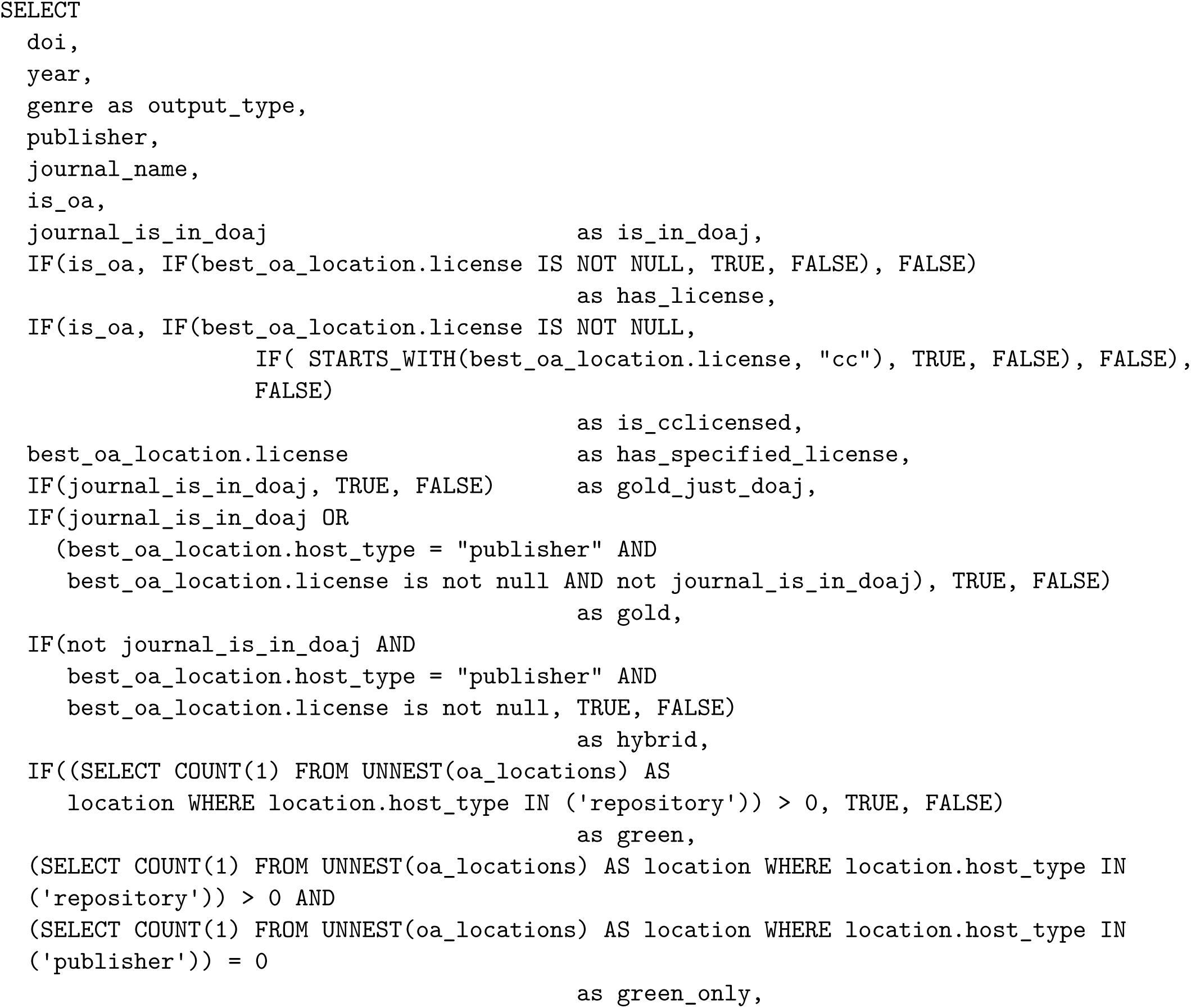
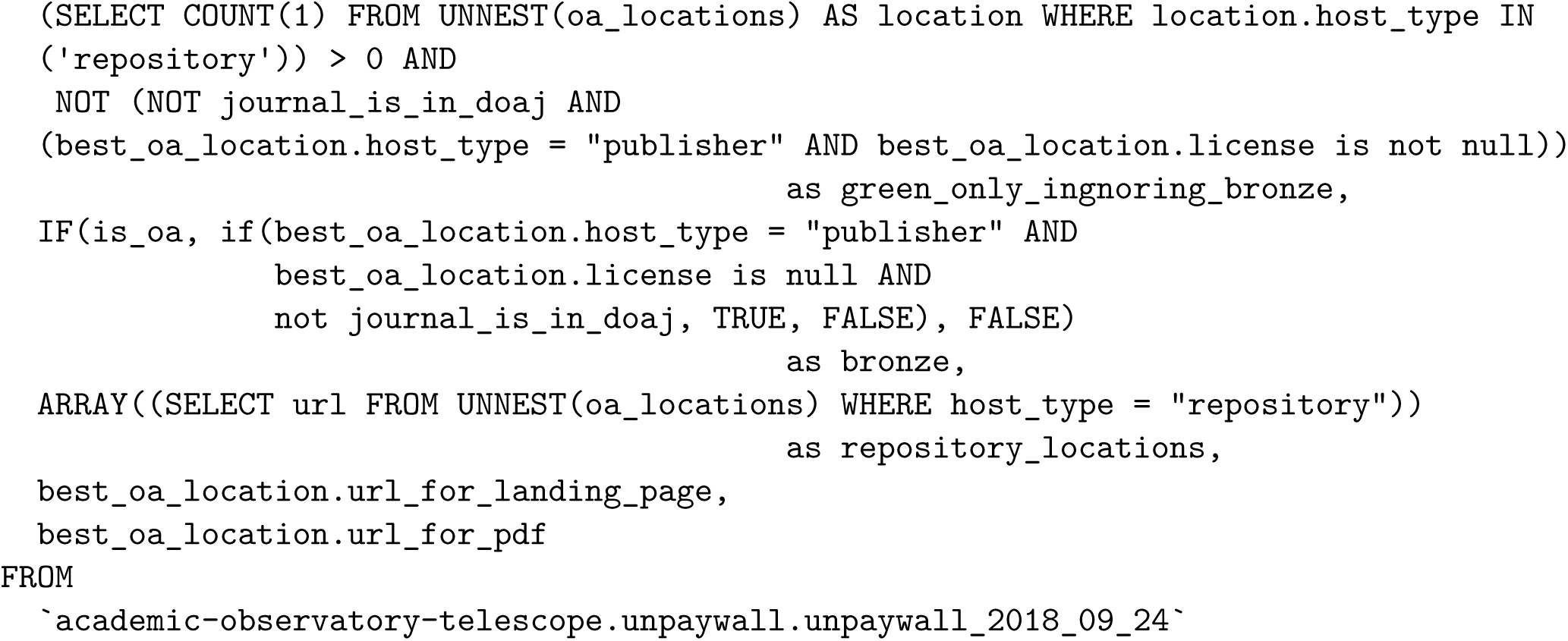

#### 3.3 Identification of grids in scope

Our full dataset includes all those institutions for which a GRID is recorded in Microsoft Academic Graph. In this article we have focussed on a set of institutions seeded from the top 1000 institutions in the THE World University Ranking supplemented for greater geographical coverage and deeper coverage of specific countries. We identify those GRIDs in scope for this article by identifying GRIDs for which we have additionally collected data from Scopus and Web of Science. As there are a small number of non-university research institutions in this set we explicitly exclude them.

The query that generates the `grids_in_scope` table is as follows:

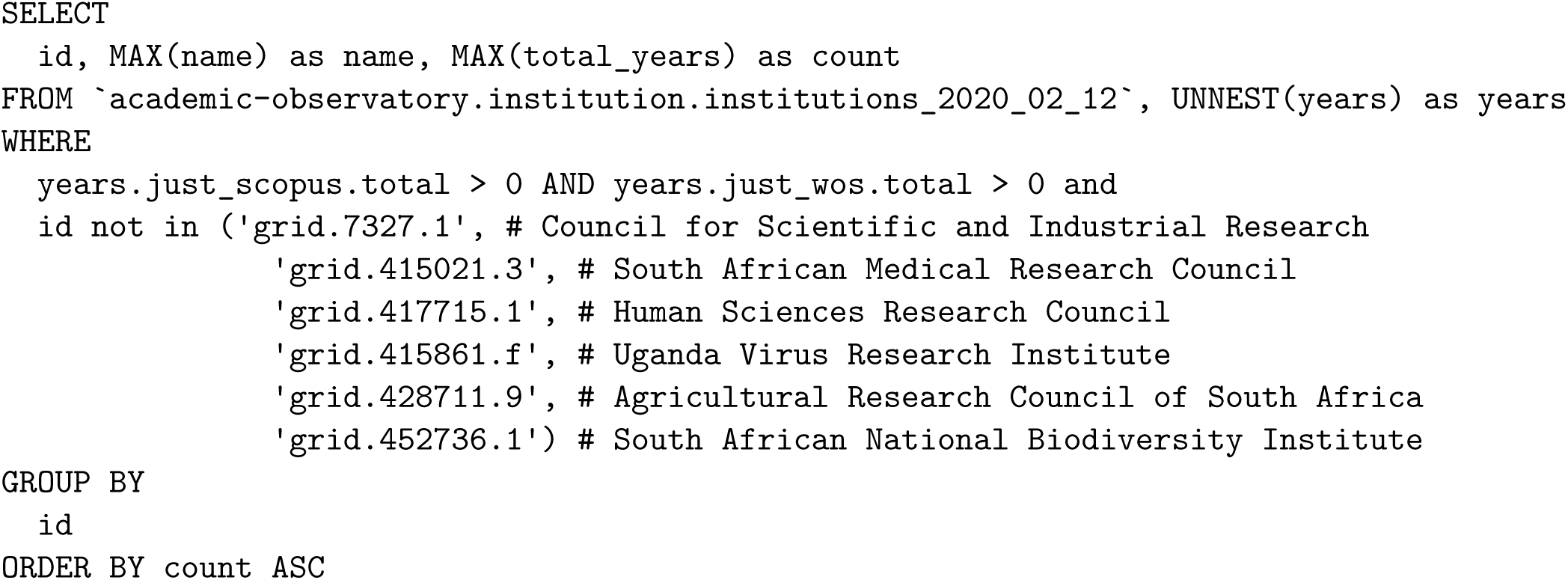

#### 3.4 Identification of named universities in the top 100s

The dataset containing named universities are all of those which fall, for any year from 2013-2018 inclusive, into the top 100 of:

1. Overall percentage of OA (i.e., Total OA)
2. Percentage of Green OA
3. Percentage of Gold OA

This group is identified using the following query and code, which is used to generate the table ‘named_institutions_in_scope’ which is also provided in the shared dataset. We select 110 from each category due to the filtering downstream of smaller institutions.

This table is generated by a small python script as follows:

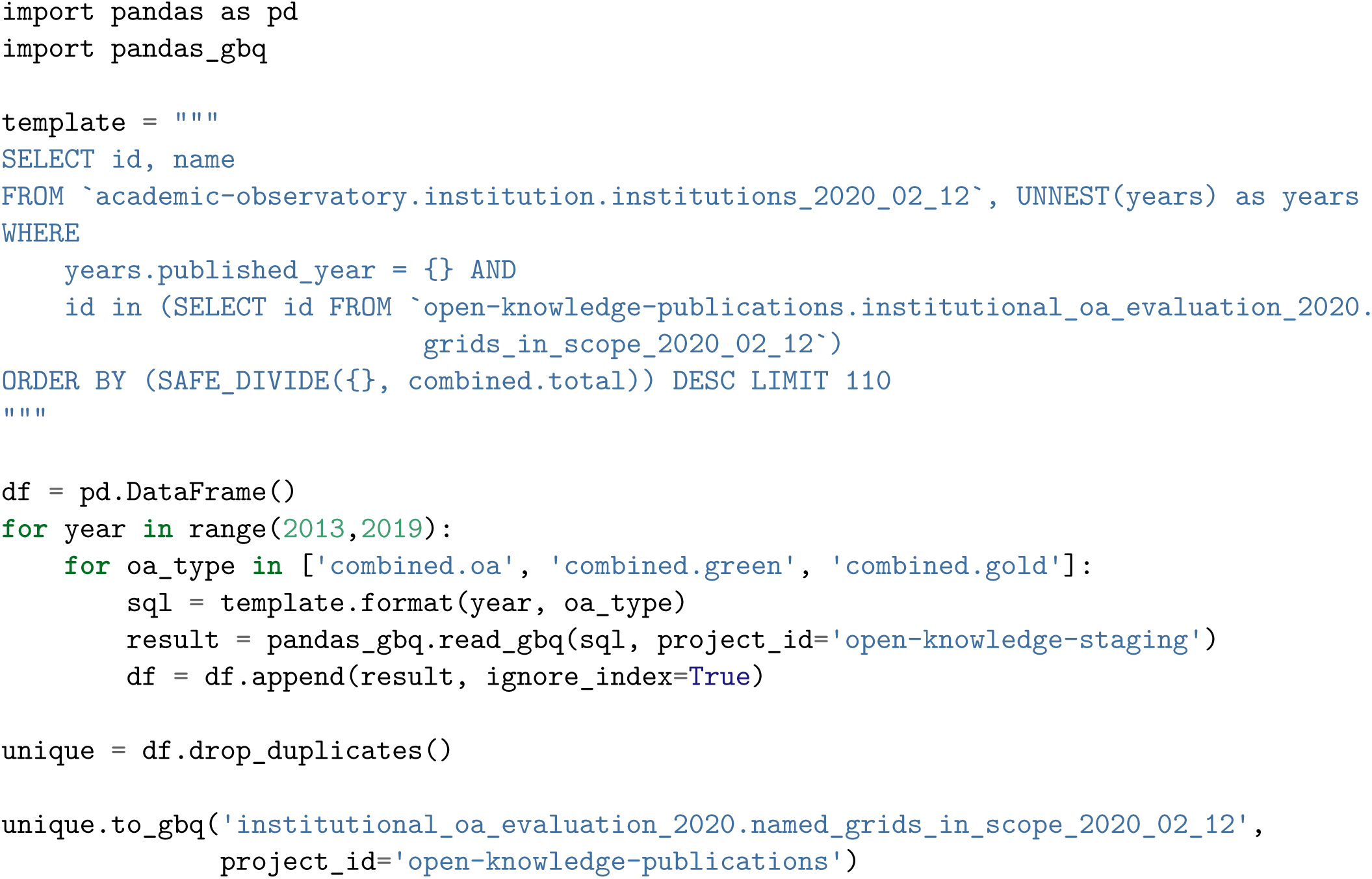

#### 3.5 Generation of the derived datasets

The four main datasets that are publicly shared are generated directly from the `institutions` table using either the `grids_in_scope` or `named_grids_in_scope` tables to provide a filter for the set of institutions. The queries have some minor differences to provide the data of interest in each case. All four queries are provided in the Data and Queries package available at Zenodo. Here we show the query for the generation of the `full_paper_dataset`. We use a salt to generate the anonymised IDs for each university.

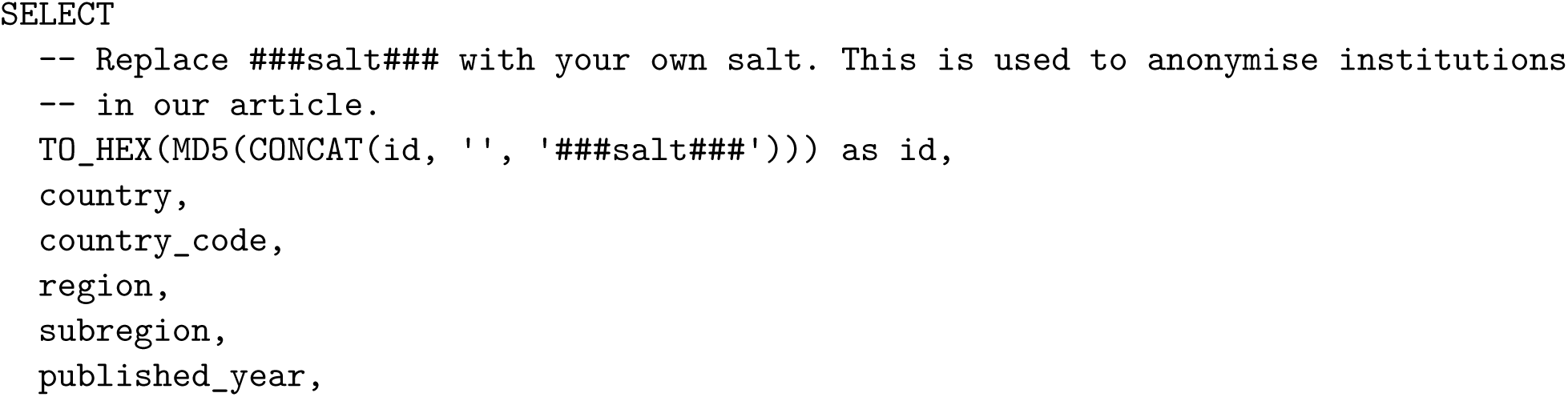
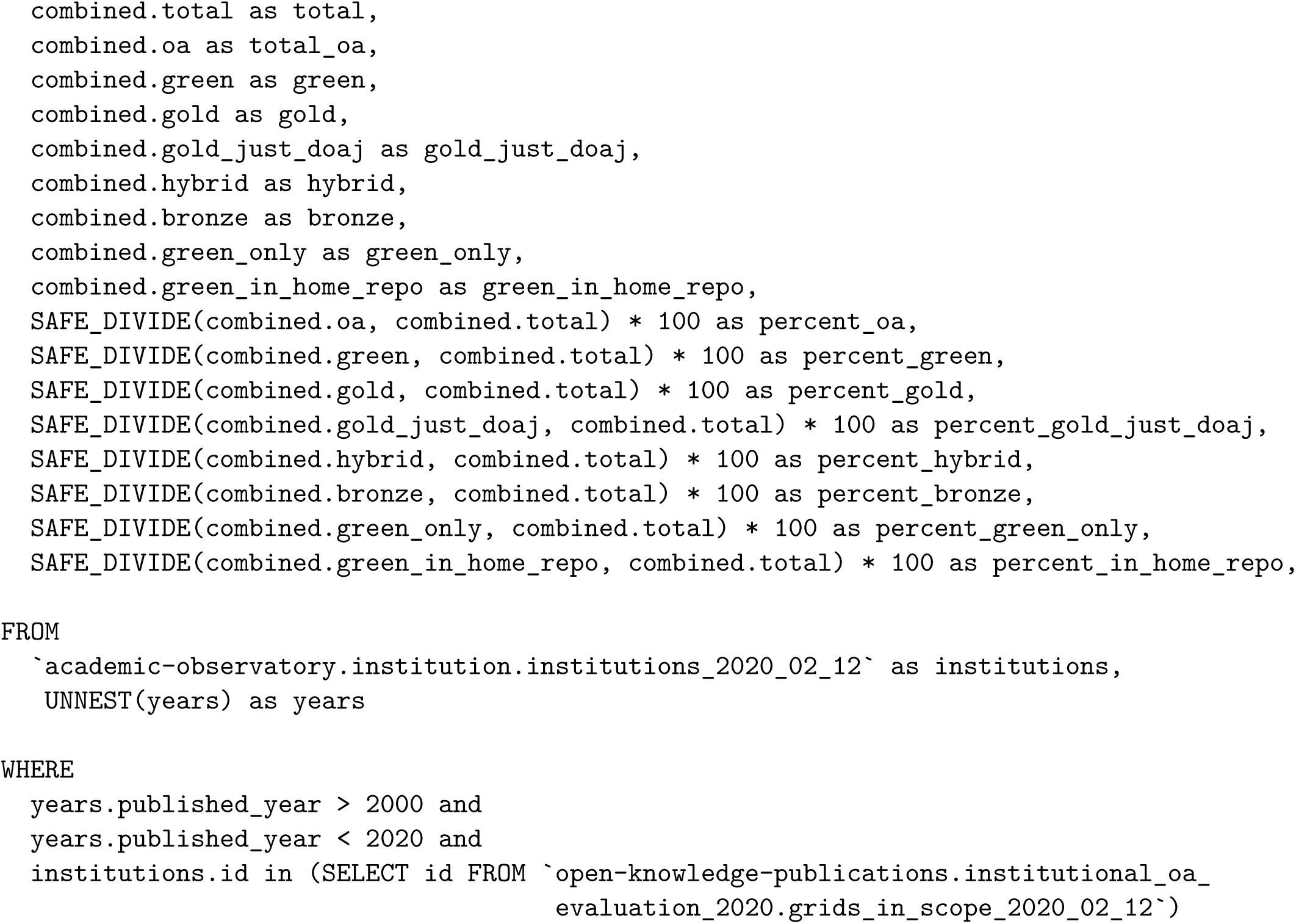

## Supplementary Figures

This section provides additional supplementary figures that are largely in parallel with those presented in the main article, apart for different publication years. Supplementary Figures 1 and 2 present the top 100 universities in total open access, open access (gold) publishing and repository-mediated (green) open access, for the publication years 2016 and 2018, respectively. These are produced analogously to Figure 1 of the main text.

Supplementary Figures 3 to 5 are figures based on the complete list of universities (i.e., without filtering on margin of error). These represent the open access performances across different countries for the years 2016, 2017 and 2018. In parallel, Supplementary Figures 6 to 8 gives the overviews as per region.

Supplementary Figure 9 is an extended version of Figure 3 of the main text and presents corresponsing time-stamped figures from 2007 to 2018. Supplementary Figure 10 provides additional information to Figure 4 of the main text.

**Supplementary Figure 1:**
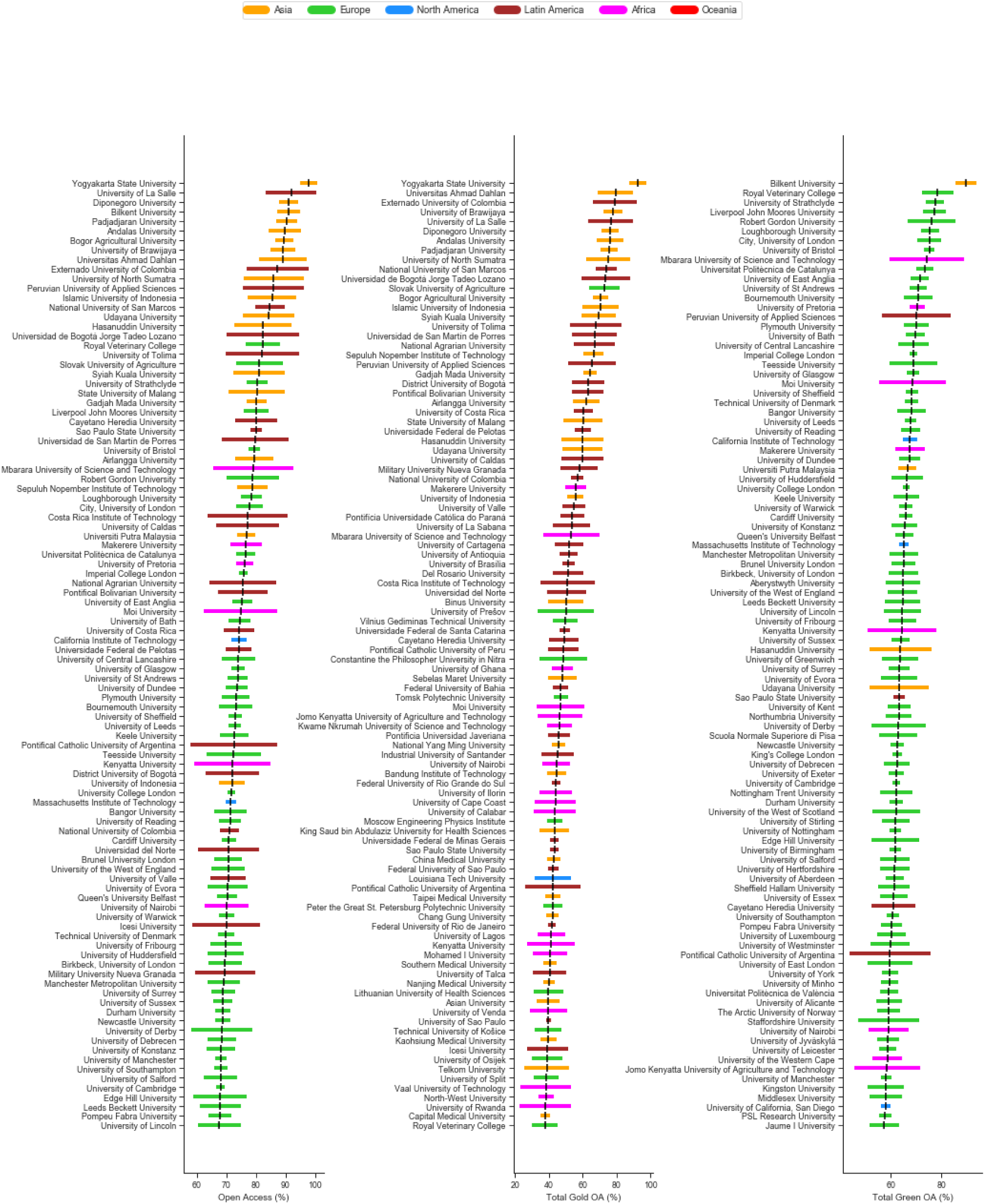
Top 100 universities in terms of performance in Total OA, Gold OA and Green OA for 2016.

**Supplementary Figure 2:**
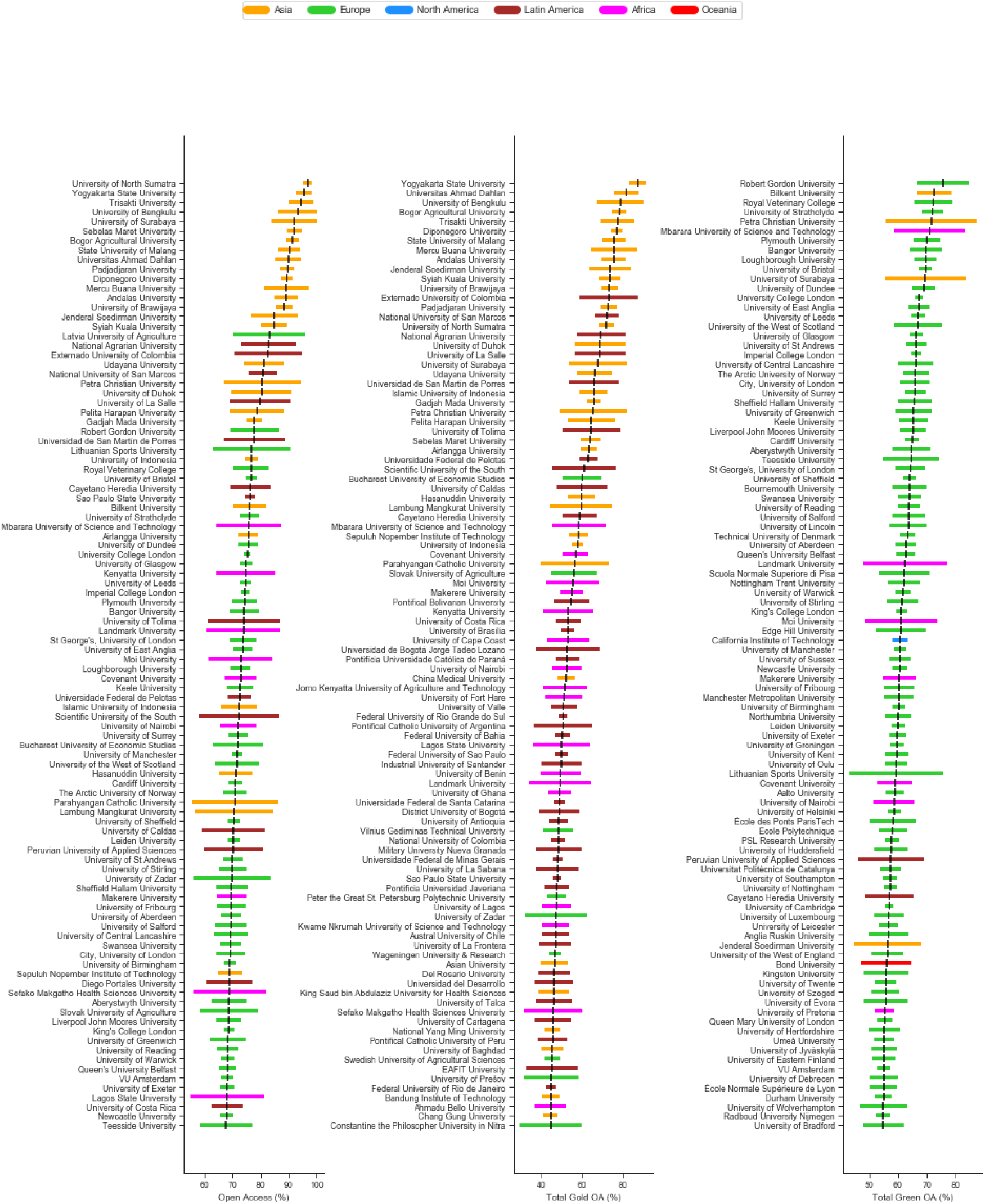
Top 100 universities in terms of performance in Total OA, Gold OA and Green OA for 2018.

**Supplementary Figure 3:**
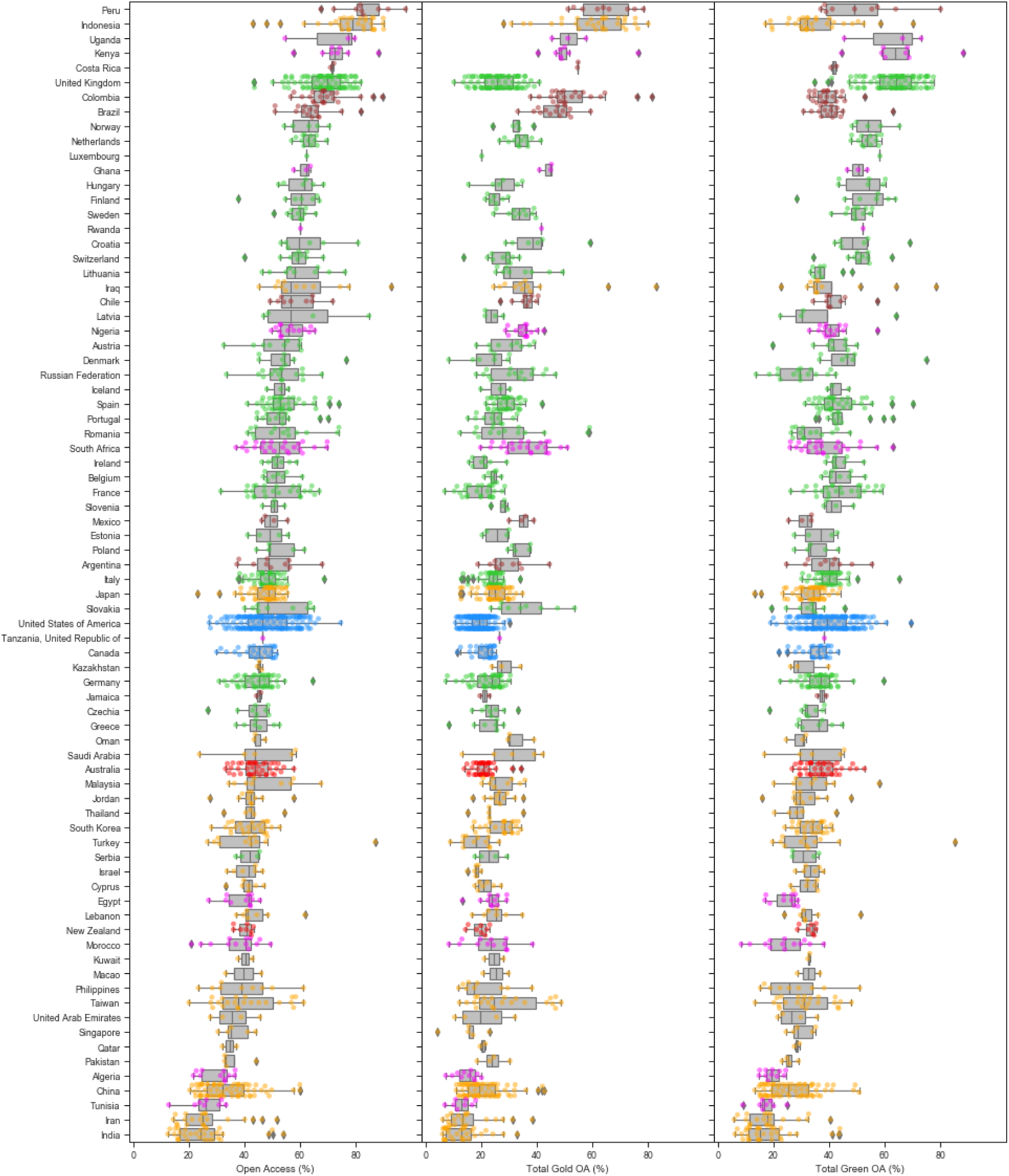
Percentages of institutional Total OA, Gold OA and Green OA (left to right) grouped by countries for 2017.

**Supplementary Figure 4:**
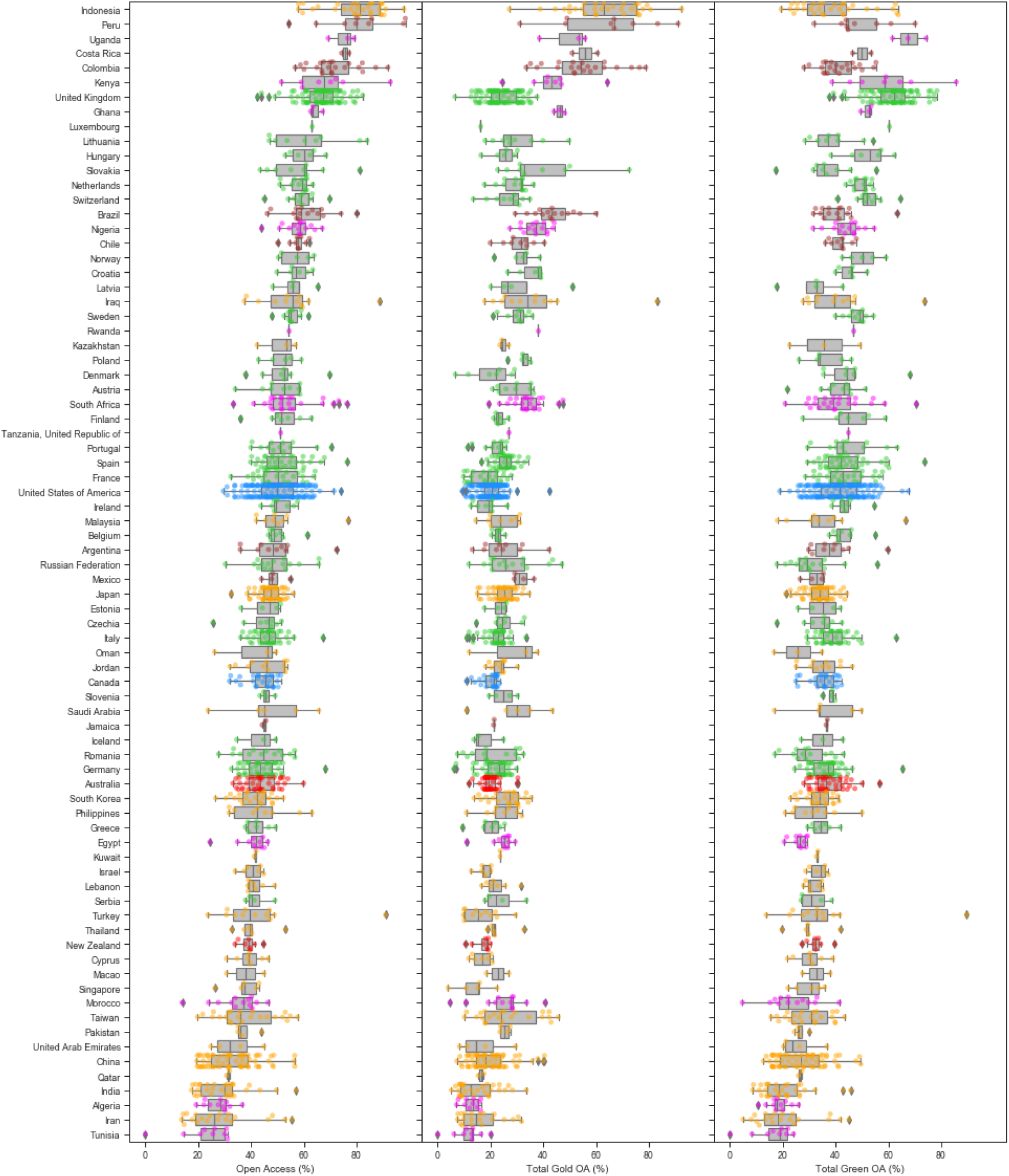
Percentages of institutional Total OA, Gold OA and Green OA (left to right) grouped by countries for 2016.

**Supplementary Figure 5:**
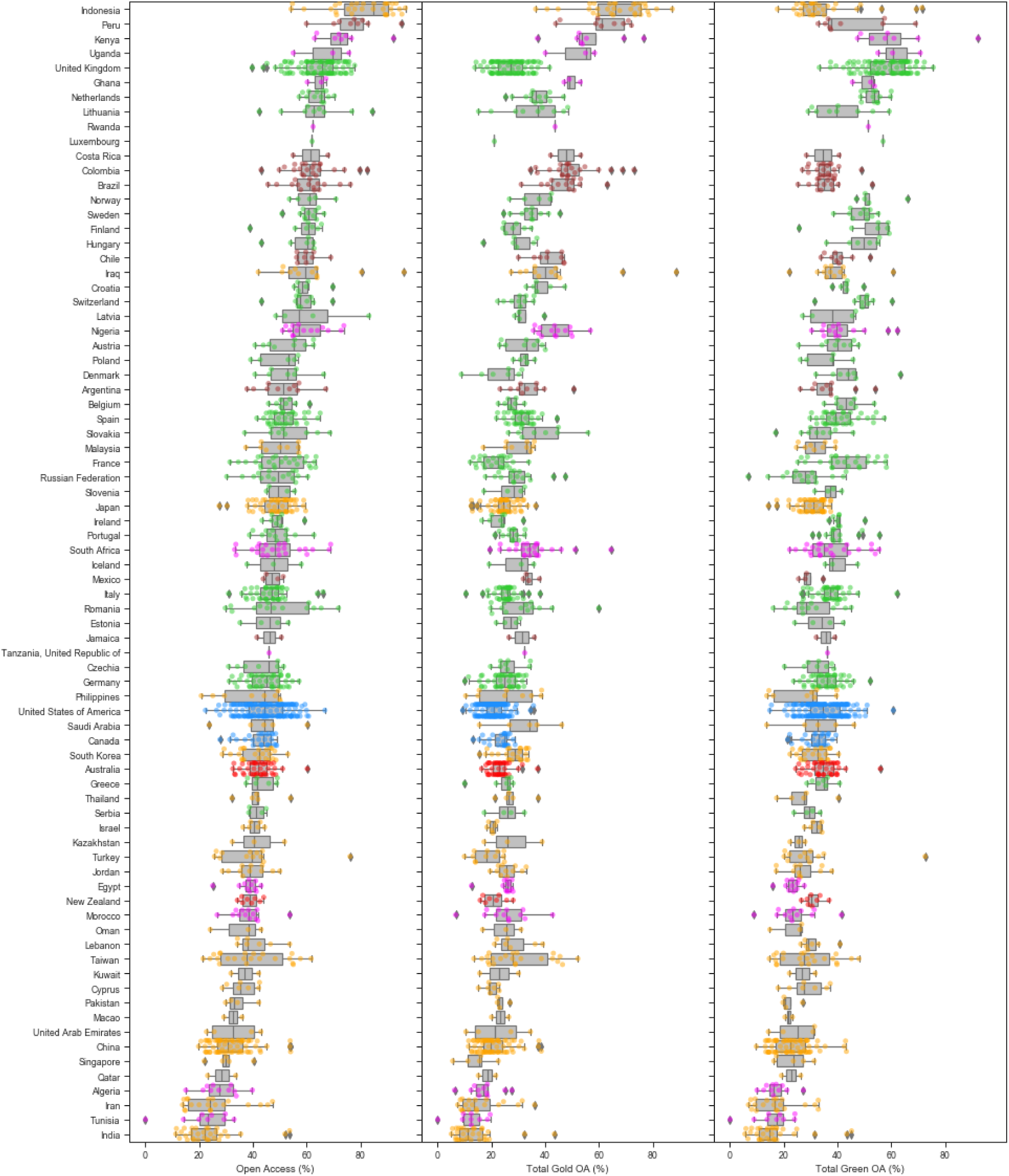
Percentages of institutional Total OA, Gold OA and Green OA (left to right) grouped by countries for 2018.

**Supplementary Figure 6:**
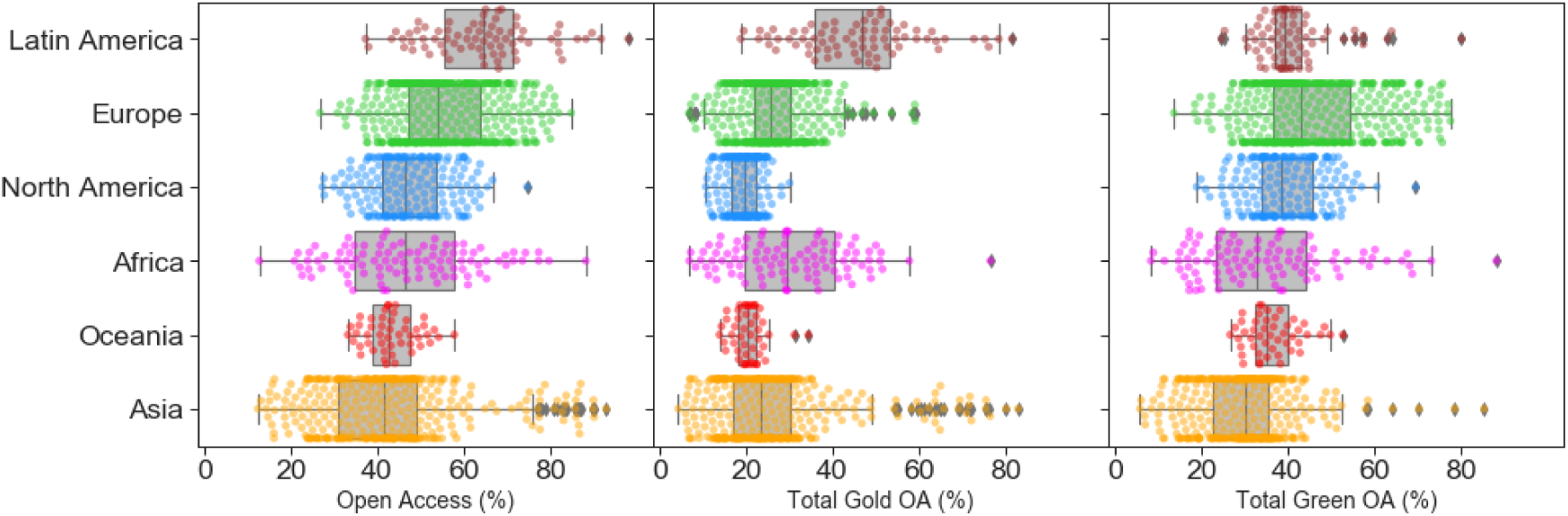
Percentages of institutional Total OA, Gold OA and Green OA (left to right) grouped by regions for 2017.

**Supplementary Figure 7:**
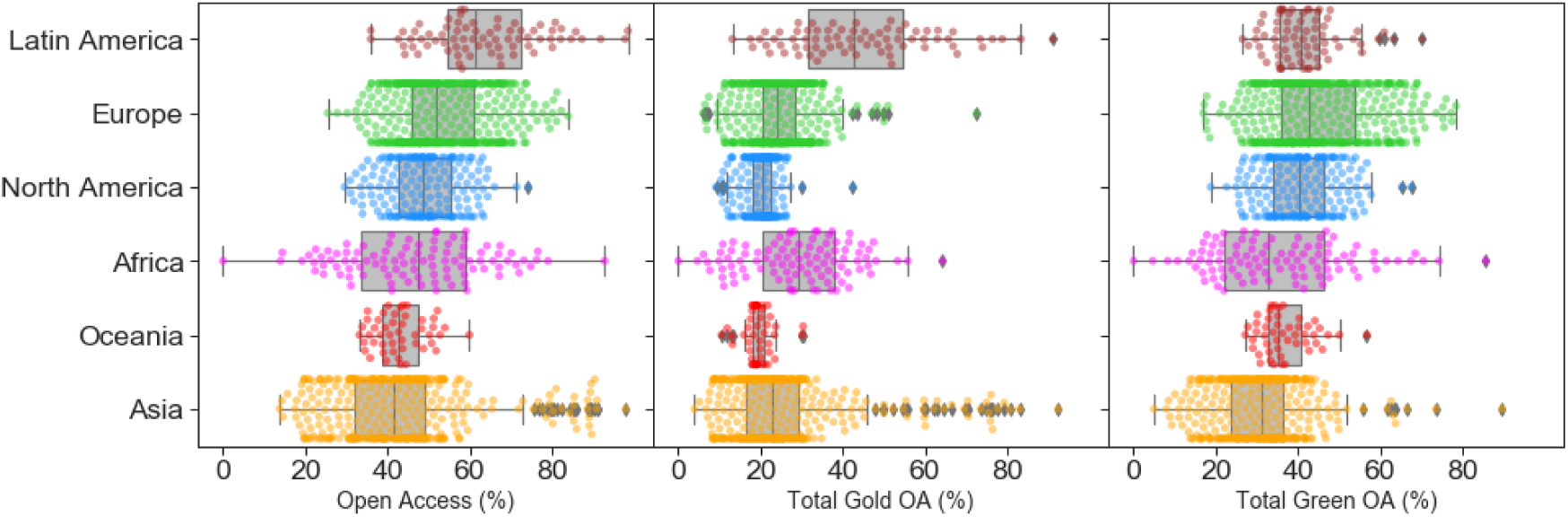
Percentages of institutional Total OA, Gold OA and Green OA (left to right) grouped by regions for 2018.

**Supplementary Figure 8:**
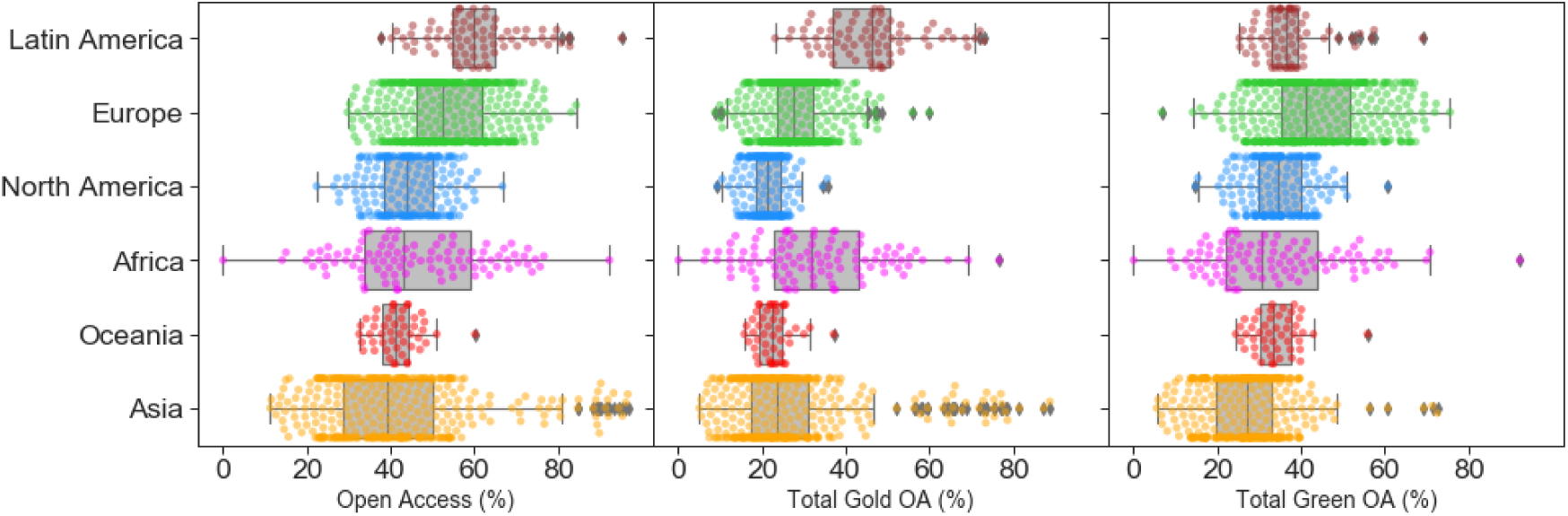
Percentages of institutional Total OA, Gold OA and Green OA (left to right) grouped by regions for 2016.

**Supplementary Figure 9:**
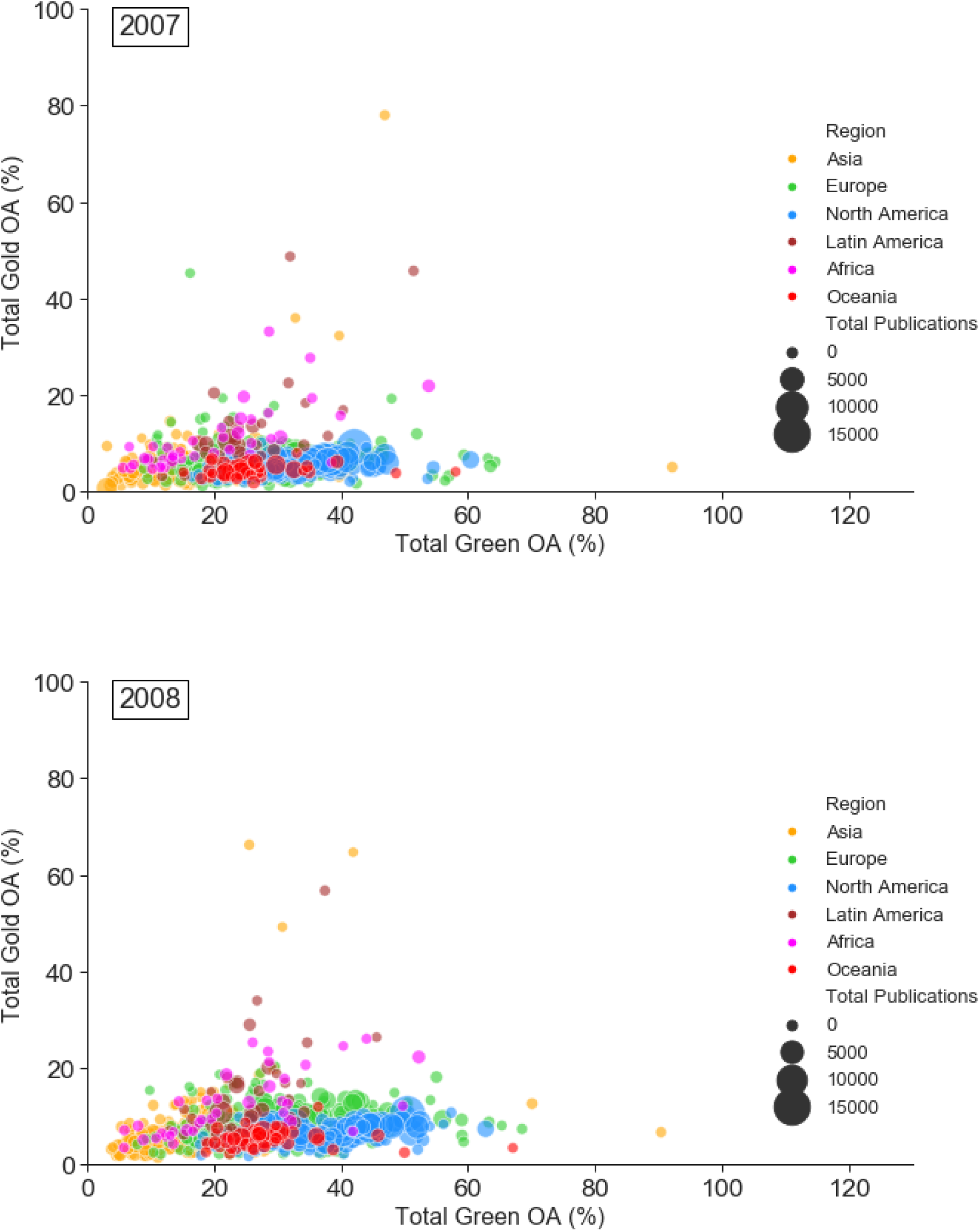
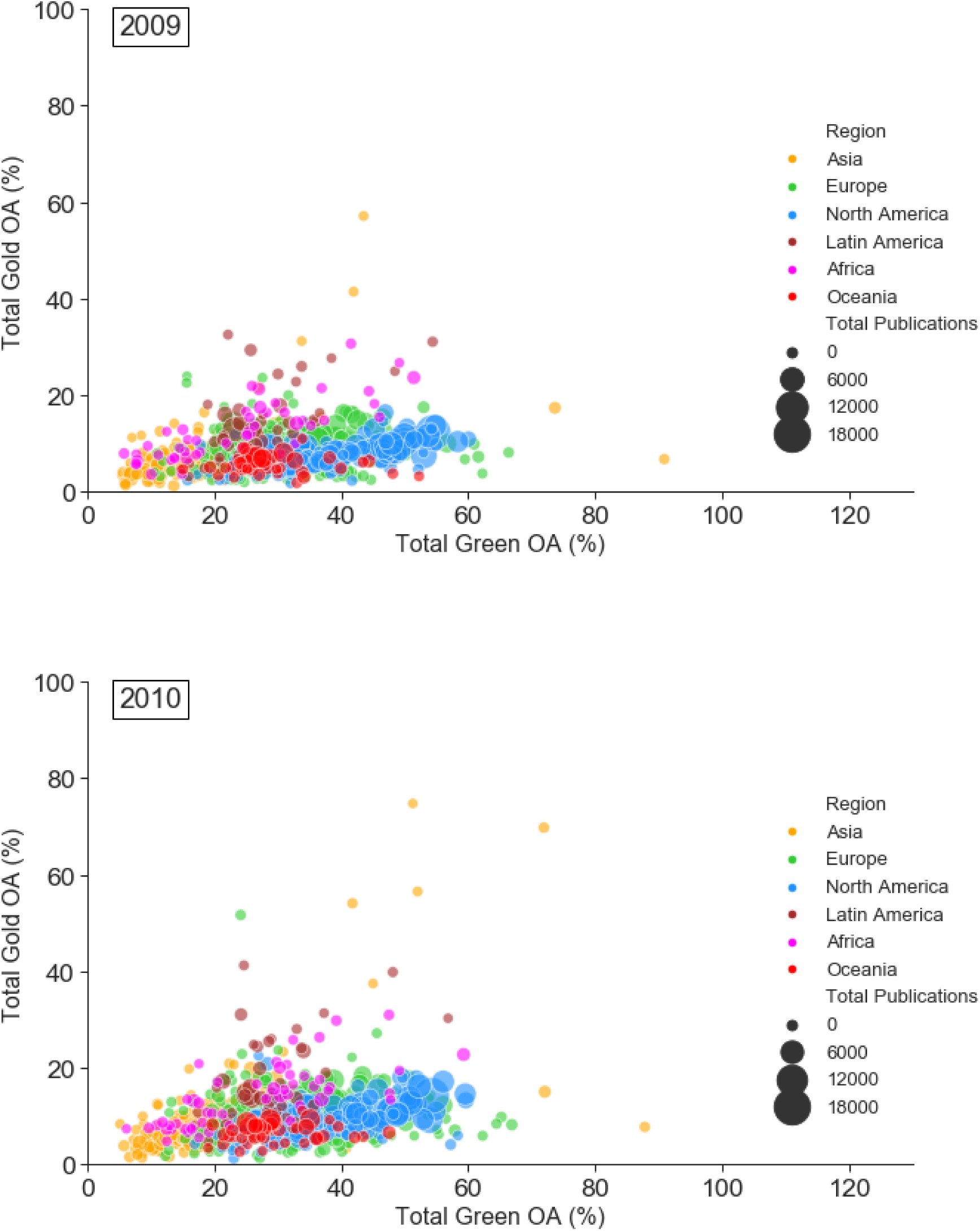
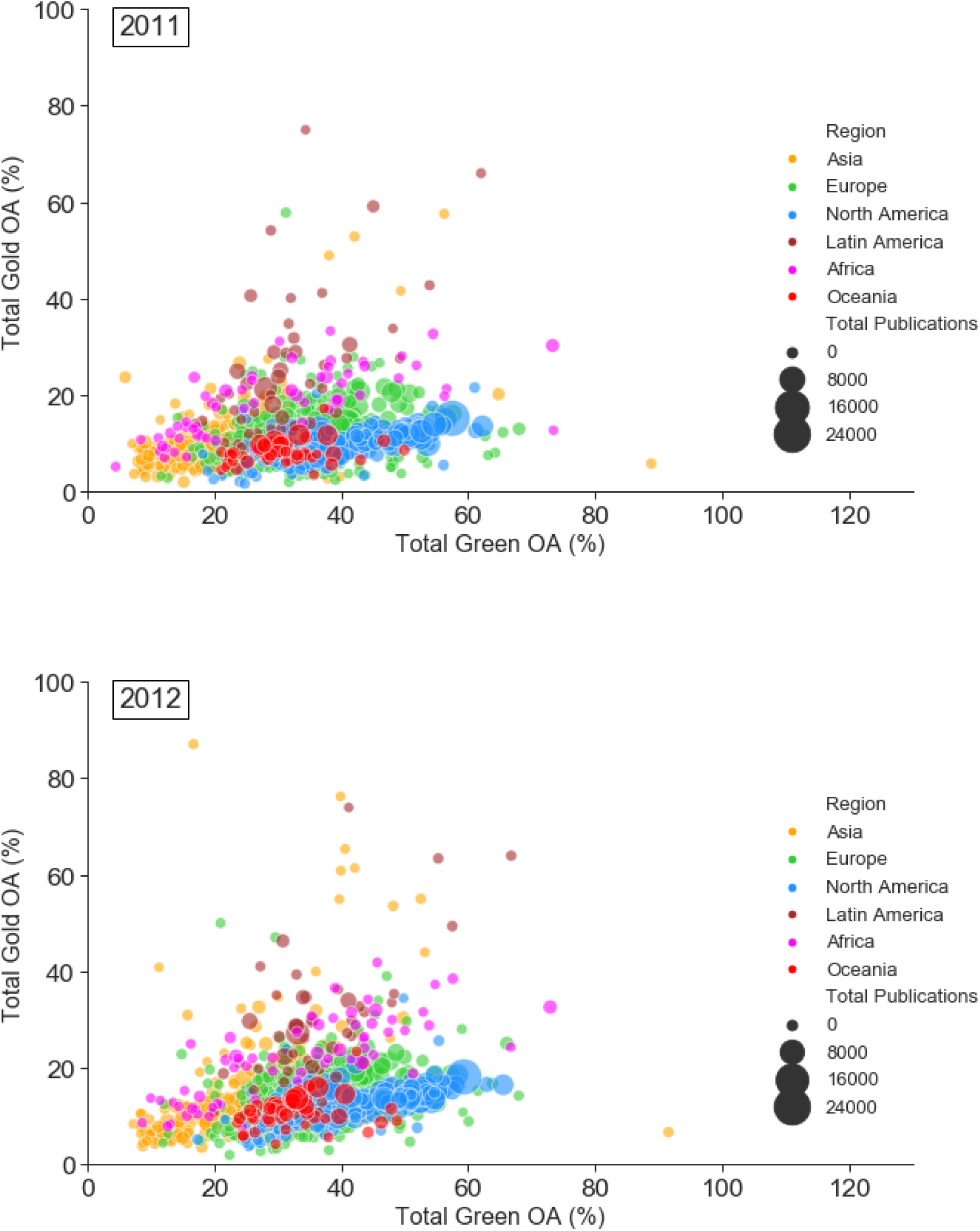
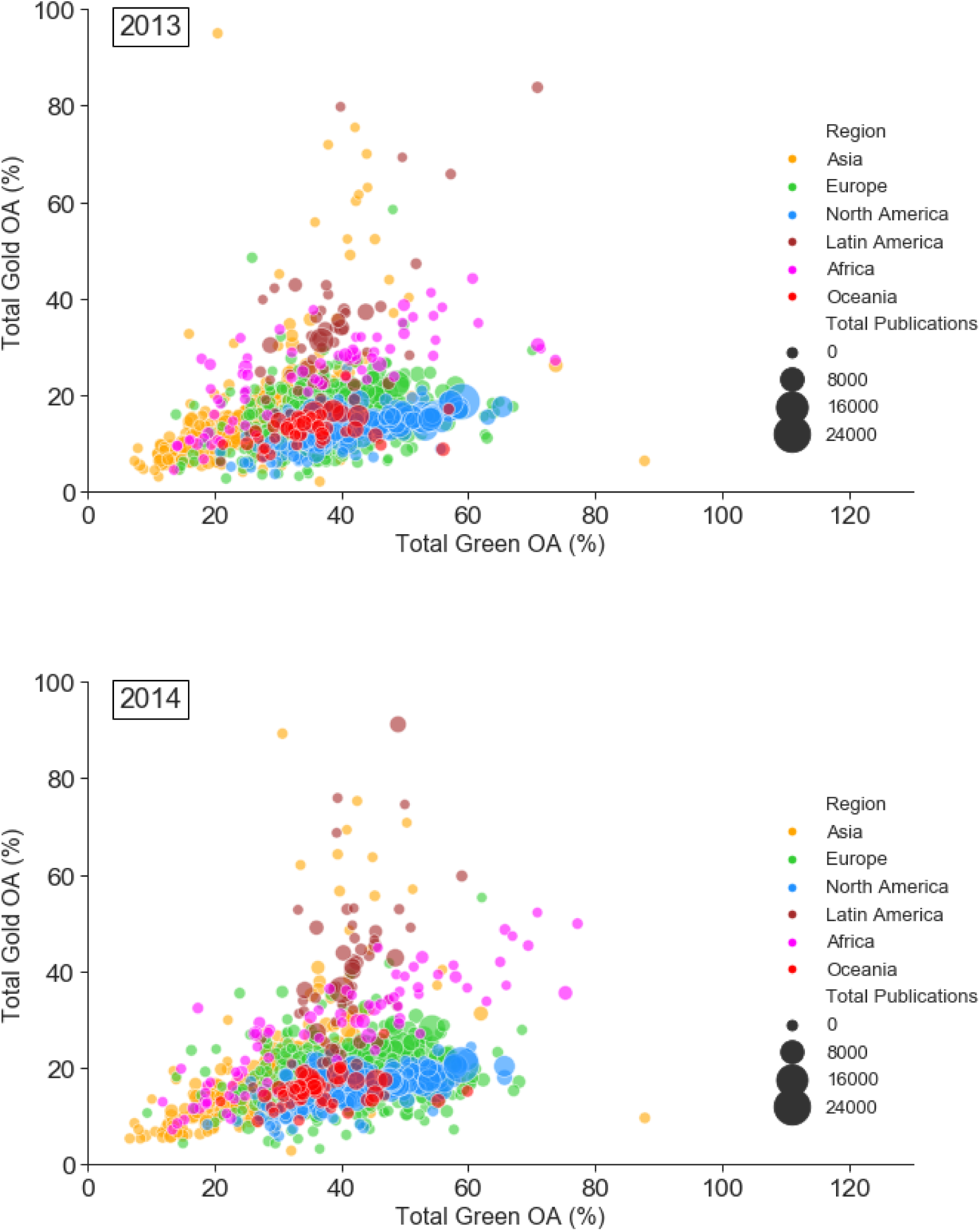
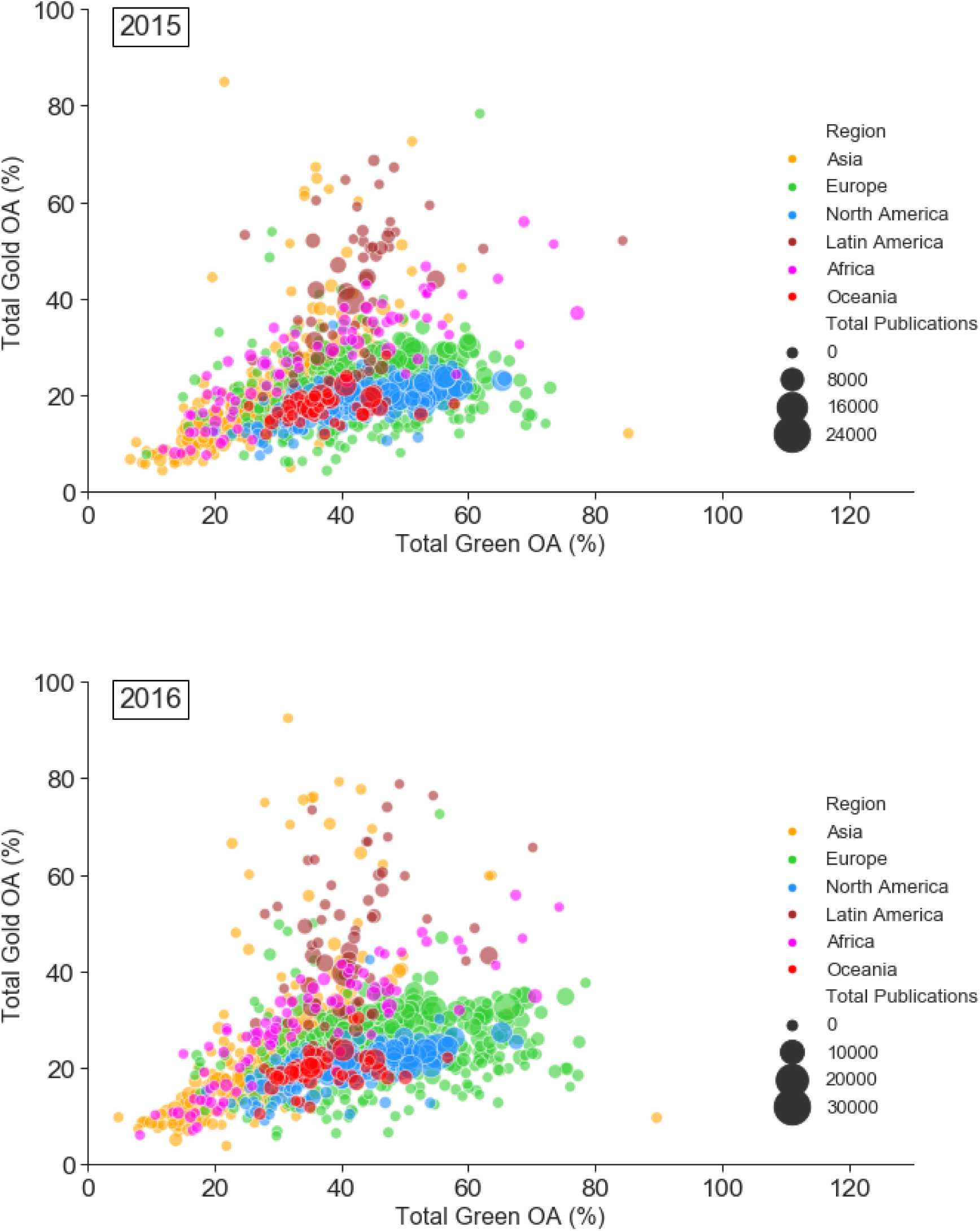
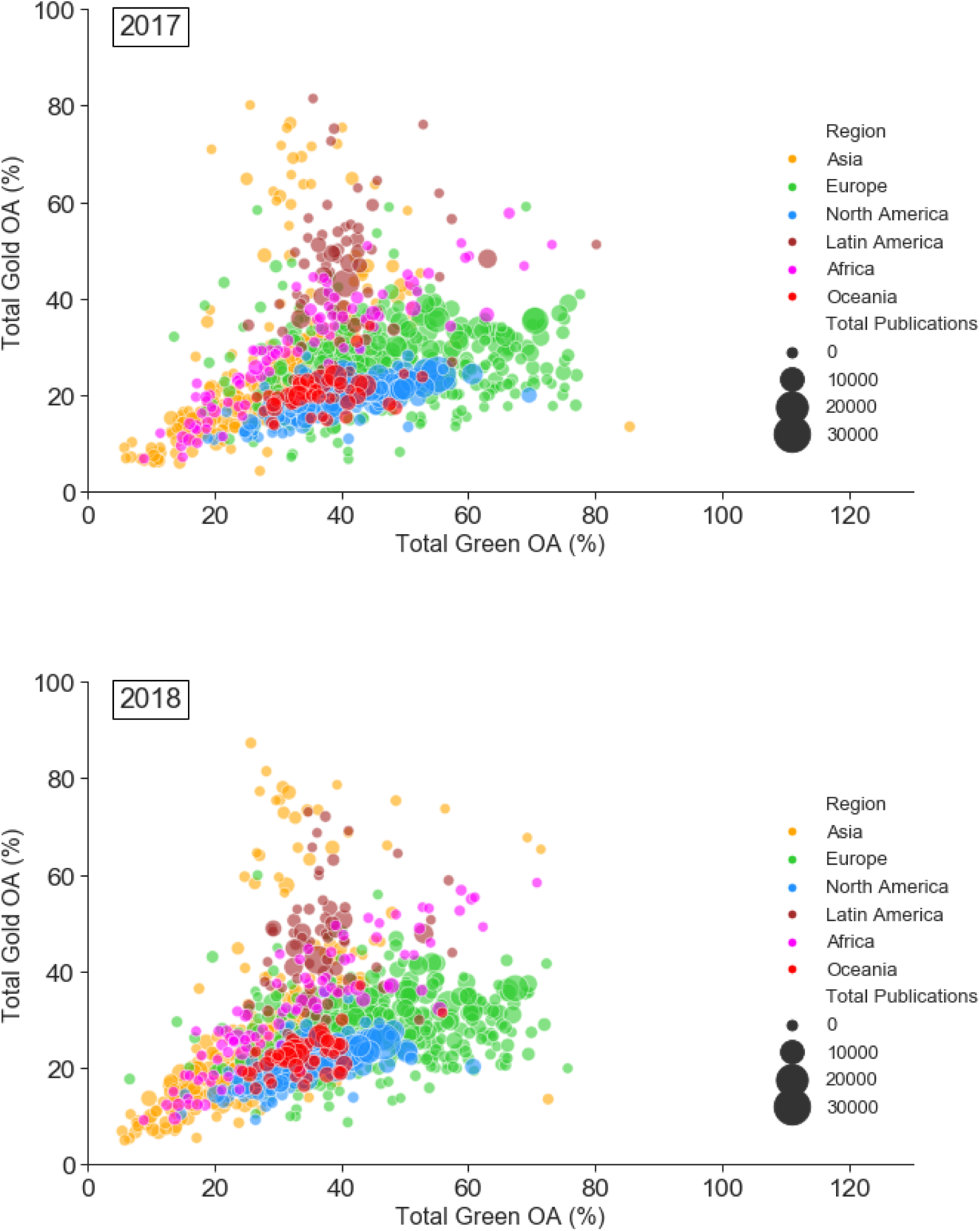
Open access publishing vs repository-mediated open access by institution from 2007-2018.

**Supplementary Figure 10:**
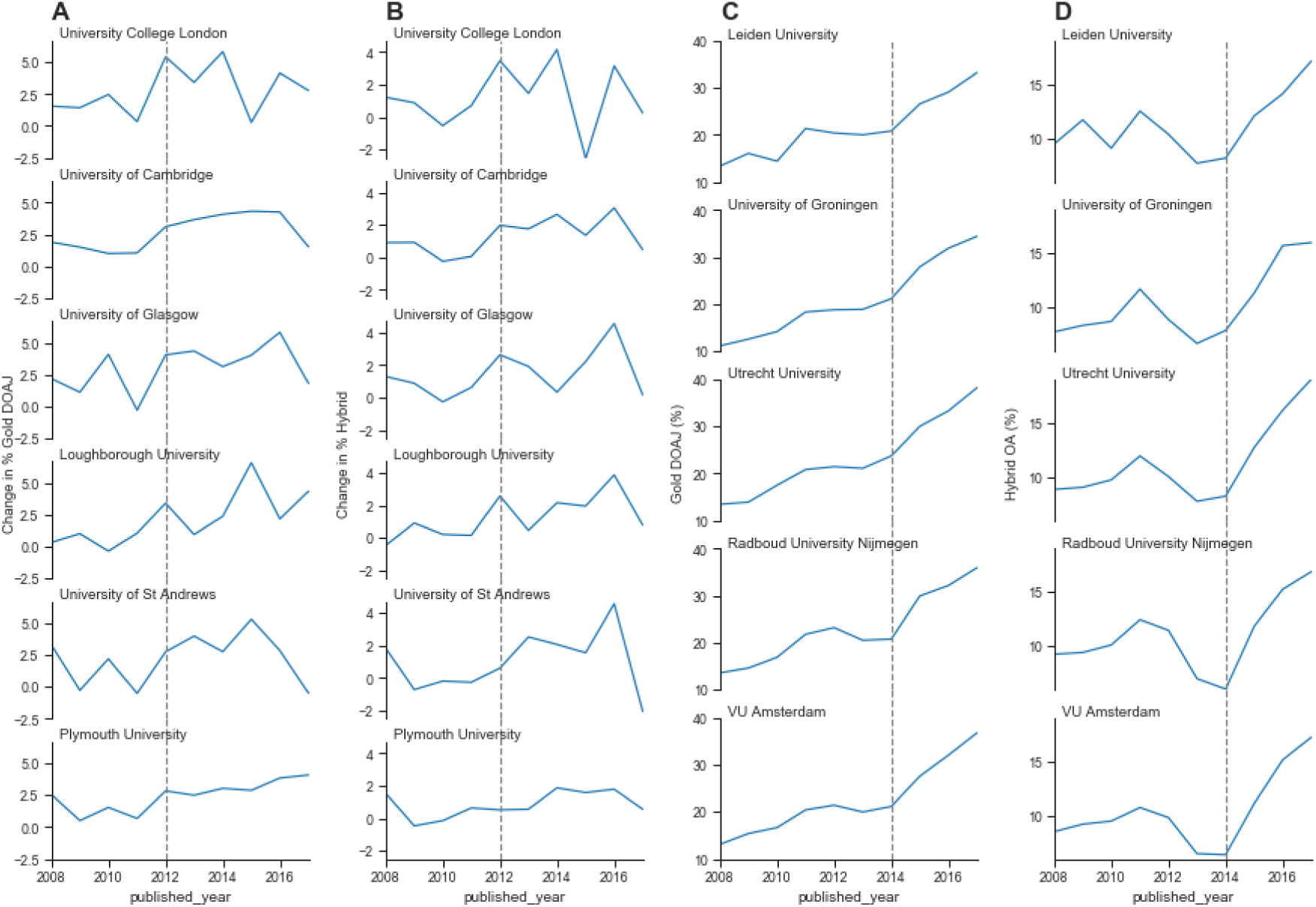
Additional figures for monitoring the effect of policy interventions for selected groups of universities.

## Footnotes

1 See https://www.coalition-s.org/

2 See https://unpaywall.org/

3 See https://aka.ms/msracad

